# Hardwired to attack: Transcriptionally defined amygdala subpopulations play distinct roles in innate social behaviors

**DOI:** 10.1101/2023.03.16.532692

**Authors:** Julieta E. Lischinsky, Luping Yin, Chenxi Shi, Nandkishore Prakash, Jared Burke, Govind Shekaran, Maria Grba, Joshua G. Corbin, Dayu Lin

## Abstract

Social behaviors are innate and supported by dedicated neural circuits, but it remains unclear whether these circuits are developmentally hardwired or established through social experience. Here, we revealed distinct response patterns and functions in social behavior of medial amygdala (MeA) cells originating from two embryonically parcellated developmental lineages. MeA cells in male mice that express the transcription factor Foxp2 (MeA^Foxp2^) are specialized for processing male conspecific cues even before puberty and are essential for adult inter-male aggression. In contrast, MeA cells derived from the *Dbx1*-lineage (MeA^Dbx1^) respond broadly to social cues and are non-essential for male aggression. Furthermore, MeA^Foxp2^ and MeA^Dbx1^ cells show differential anatomical and functional connectivity. Altogether, our results support a developmentally hardwired aggression circuit at the level of the MeA and we propose a lineage-based circuit organization by which a cell’s embryonic transcription factor profile determines its social information representation and behavior relevance during adulthood.

**Highlights:**

- MeA^Foxp2^ cells in male mice show highly specific responses to male conspecific cues and during attack while MeA^Dbx1^ cells are broadly tuned to social cues.
- The male-specific response of MeA^Foxp2^ cells is present in naïve adult males and adult social experience refines the response by increasing its trial-to-trial reliability and temporal precision.
- MeA^Foxp2^ cells show biased response to males even before puberty.
- Activation of MeA^Foxp2^, but not MeA^Dbx1^, cells promote inter-male aggression in naïve male mice.
- Inactivation of MeA^Foxp2^, but not MeA^Dbx1^, cells suppresses inter-male aggression.
- MeA^Foxp2^ and MeA^Dbx1^ cells show differential connectivity at both the input and output levels.

## Introduction

Innate social behaviors, such as mating, fighting and parenting, are indispensable for the survival and propagation of a species, and therefore present widely in the animal kingdom. These behaviors are considered innate as they can take place without learning although the efficiency in performing these behaviors can be improved with repeated execution^1^. The developmental mechanisms for the establishment of innate social behaviors and the role of experience in shaping these circuits remain poorly understood.

An array of interconnected brain regions, collectively referred to as the social behavior network (SBN), were proposed to be important for diverse social behaviors^2,3^. The medial amygdala (MeA), especially its posterior division (pMeA), is considered a key node of the SBN based on its connectivity, activity, gonadal hormone receptor expression and numerous lesion studies^2^. At the input level, MeA is the primary recipient of accessory olfactory bulb (AOB) inputs-the exclusive relay of the vomeronasal organ (VNO) specialized in detecting pheromones^4^. Volatile information from the main olfactory system also converges onto MeA cells via the cortical amygdala^5,6^. Consistent with the strong olfactory inputs, c-Fos, a surrogate of neural activation, is highly expressed in the MeA following exposure to conspecific chemosensory cues^7-9^. *In vivo* electrophysiological recording and Ca*^2+^* imaging further revealed response of MeA cells to a wide array of conspecific and heterospecific cues, including males, females, pups, and predator odors^6,10^. Unsurprisingly, MeA lesion, which impedes the flow of social sensory information, causes deficits in multiple social behaviors, including male sexual behavior, aggression and maternal behaviors^11-15^. These studies collectively support an important role for the MeA in processing and relaying olfactory information related to conspecifics.

Recent functional experiments suggest a more direct role of pMeA in driving social behaviors. Hong et. al. first showed that optogenetic activation of GABAergic pMeA cells (the major cell type in the dorsal pMeA) acutely induced mounting or attack in male mice depending on stimulation intensity^16^. Later, Unger et. al. reported that silencing or ablating aromatase expressing MeA cells decreased aggression in both males and females^17^. Padilla et. al. found that optogenetic activation of the projection from MeA Npy1r expressing cells to bed nucleus of the stria terminalis (BNST) promoted male aggression^18^ and Miller et. al. reported similar aggression-promoting effect of the MeA^D1R^ to BNST pathway^19^. Nordman et. al. showed that high frequency stimulation of MeA CaMKII cells could prime aggression through its projection to BNST and ventromedial hypothalamus (VMH)^20^. Most recently, MeA GABAergic cells were also found to drive social behaviors besides aggression, including pup grooming, infanticide and allogrooming^21,22^.

These results raised several questions regarding the MeA function in social behaviors. First, does the MeA mainly encode olfactory cues or also carry action-related information? While MeA cells have been consistently found to be activated by conspecific olfactory cues^6,7,10^, the responses of MeA cells during the action phase of adult-directed social behaviors, such as attack and mount, remains unexplored. Second, are there dedicated subpopulations in the MeA for distinct social behaviors or can any random subsets of MeA cells generate any social behavior in a context and intensity dependent manner? An answer to this question remains unclear as activating multiple subpopulations of MeA cells can all induce aggression^17-20^, while activating the same GABAergic MeA population induces diverse social behaviors^16,21,22^. Third, how much of the MeA cell response is developmentally hardwired vs. determined by adult experience? Through immediate-early gene mapping, Choi et. al. found that MeA cells relevant for social behaviors and predator defense are marked by distinct transcription factors, suggesting developmental hardwiring of social vs. non-social signals^7^. However, recent imaging studies revealed that MeA cell responses to social stimuli can be altered with adult experience, suggesting that the exact social response of MeA cells may not be pre-determined^23^.

Taken together, although the MeA is clearly a central node of SBN, how the MeA mediates social behaviors remains elusive. In our previous studies, we identified two distinct MeA populations that arise from separate embryonic lineages in the telencephalic preoptic area (POA), marked by the transcription factors, Dbx1 and Foxp2^9,24^. In adults, although Dbx1 is no longer expressed in the MeA, *Dbx1*-lineage cells remain distinct from Foxp2 expressing cells despite being spatially intermingled^9^ (**Figure 1**). In addition, these two subpopulations differ in their gene expression patterns and intrinsic electrophysiological properties^9^. Therefore, we reason that these two developmentally distinct and transcriptionally-defined subpopulations could provide a unique opportunity to address whether MeA cells are hardwired for social behaviors or not. Specifically, are social cue representation and social function of MeA cells predetermined by their developmental lineage? Here, we performed a series of *in vivo* population recordings, functional manipulations and tracing experiments to compare the neuronal responses, functions, and connectivity of MeA^Dbx1^ and MeA^Foxp2^ cells in male social behaviors and revealed the response pattern of MeA^Foxp2^ cells over development.

**Figure 1.**
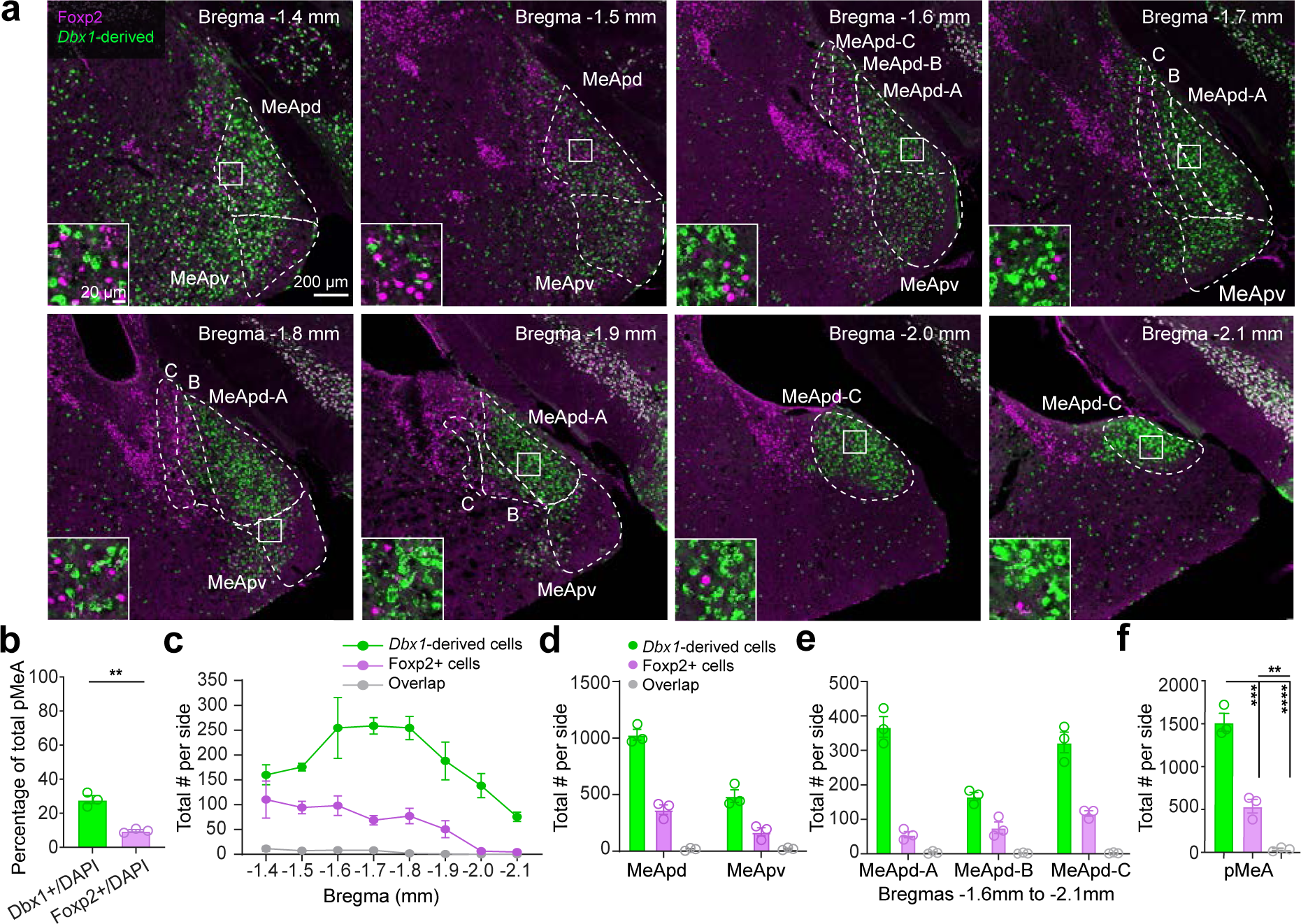
MeA^Foxp2^ and MeA^Dbx1^ cells are non-overlapping transcriptionally defined subpopulations. **(a)** Immunostaining of Foxp2 and GFP (*Dbx1*-derived cells) in the MeA of Dbx1^cre^;Ai6 male mice. Left bottom shows the enlarged view of boxed areas. **(b)** Percentage of MeA^Foxp2^ and MeA^Dbx1^ cells in the total MeA population. **(c)** The number of counted Foxp2, *Dbx1*-derived and double positive cells in each side of the MeA from Bregma −1.4mm to −2.1mm. **(d)** The total number of counted Foxp2, *Dbx1*-derived and double positive cells in each side of the posterodorsal and posteroventral MeA (MeApd and MeApv). **(e)** The total number of counted Foxp2, *Dbx1*-derived and double positive cells in the MeApd sub-compartments from Bregma −1.6mm to −2.1mm. **(f)** Total number of Foxp2, *Dbx1*-derived and overlap cells in each side of the posterior MeA. For b-f, every third of 50µm brain sections were counted. The Allen Brain Reference Atlas was used to determine the MeA subdivisions and sub-compartments. (b) Two-tailed unpaired t-test. (e) One-way ANOVA followed by Tukey’s multiple comparisons test. Data are mean ± S.E.M., *p<0.05, **p<0.01, ***p<0.001, ****p<0.0001, *n=*3 mice.

## Results

### Distribution of MeA^Dbx1^ and MeA^Foxp2^ cells in male mice

To visualize the spatial distribution of MeA^Dbx1^ and MeA^Foxp2^ cells in adults, we crossed *Dbx1^cre^* mice^25^ with a ZsGreen reporter line (Ai6)^26^ and immunostained for Foxp2. MeA^Dbx1^ cells make up approximately 28% of total posterior MeA cells (MeAp, Bregma level −1.4 to −2.1mm) and are found in both dorsal and ventral subdivisions (MeApd and MeApv) (**Fig. 1a-c)**. In comparison, MeA^Foxp2^ cells are relatively fewer, constituting only 10% of pMeA cells, and largely absent from caudal MeA (**Fig. 1a-c**). Between MeApd and MeApv, both MeA^Dbx1^ and MeA^Foxp2^ cells show a dorsal bias: with approximately twice as many cells in MEApd than MeApv (**Fig.1d**). Within the MeApd, Foxp2 cells are most prominent in the lateral compartment while *Dbx1*-derived cells are biased towards the medial compartment (**Fig. 1e**). Importantly, consistent with our previous study, MeA^Dbx1^ and MeA^Foxp2^ are largely distinct, even when they occupy the same MeA region (**Fig. 1c-1f**). Of all MeA^Foxp2^ and MeA^Dbx1^ cells, only 1.8% are double positive.

### Distinct MeA^Foxp2^ and MeA^Dbx1^ cell responses to social sensory cues in head-fixed naïve male mice

To address whether MeA^Foxp2^and MeA^Dbx1^ are hardwired to respond to different social cues, we recorded the Ca^2+^ activity of each population in head-fixed naive adult male mice while presenting various social stimuli in a pseudo-random order (**Fig. 2a**). Naïve mice are animals without any social interaction except with their littermates. To record MeA^Foxp2^ cells, we injected a Cre-dependent GCaMP6f virus into the MeA of *Foxp2^cre+/-^* male mice^27^ (Foxp2^GCamP^). To record MeA^Dbx1^ cells, we generated *Dbx1^cre+/-^;LSL-FlpO^+/-^* mice. In these animals, the transient Cre expression during development leads to permanent Flp expression allowing targeting of *Dbx1*-derived cells in adult mice^28^. We injected either a Flp-dependent GCaMP6f or a Flp-dependent Cre virus together with a Cre-dependent GCaMP6f virus, into the MeA of *Dbx1^cre^;LSL-FlpO* male mice (Dbx1^GCamP^) (**Fig. 2b**). A 400-µm optic fiber was placed above the injection site to collect fluorescence signal (**Fig. 2b**). Histological analysis revealed that 88% of GCaMP6f cells express Foxp2 in Foxp2^GCamP^ mice while only 5% of GCaMP6f cells were co-labeled with Foxp2 in Dbx1^GCamP^ mice, confirming the specificity of the recorded populations (**Fig. 2c, d**).

**Figure 2.**
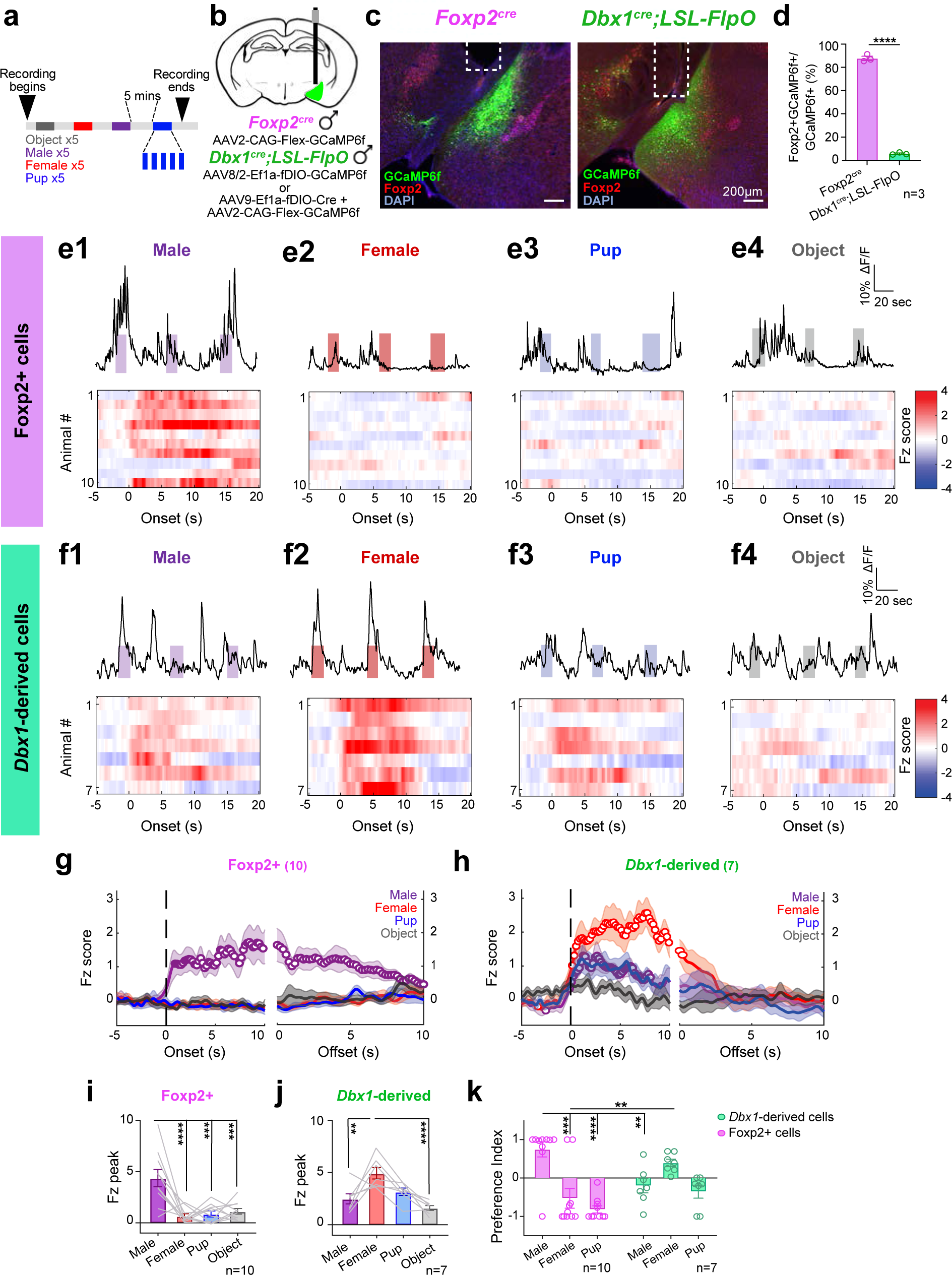
Distinct responses to social cues of MeA^Foxp2^ and MeA^Dbx1^ cells in head-fixed naïve mice. **(a)** Schematics showing the timeline of stimulus presentation. **(b)** Schematics of viral injection strategy for targeting MeA^Foxp2^ and MeA^Dbx1^ cells. **(c)** Representative histology images of viral injection, denoting GCaMP6f expression (green), Foxp2 antibody (red) and DAPI (blue) staining in Foxp2^cre^ and Dbx1^cre^;LSL-FlpO mice. White dotted lines represent location of fiber implant. **(d)** Percentage of cells co-expressing Foxp2 and GCaMP6f over the total number of GCaMP6f cells in the MeA of Foxp2^cre^ and Dbx1^cre^;LSL-FlpO mice. **(e1-e4)** Top: Representative Ca^2+^ traces of MeA^Foxp2^ cells during the presentation of a male (e1), female (e2), pup (e3) and object (e4) stimuli. Colored shades represent the duration of stimulus presentation. Bottom: corresponding heat-maps of the z-scored Ca^2+^ responses (Fz score) per animal before and after the onset of each stimuli in MeA^Foxp2^ cells. **(f1-f4)** Responses of MeA^Dbx1^ cells to various stimuli in head-fixed naïve male mice. **(g and h)** Average peri-stimulus histograms (PSTH) of Ca^2+^ signals from MeA^Foxp2^ **(g)** and MeA^Dbx1^ cells **(h)** aligned to the onset (left) and offset (right) of various stimulus presentations. Open circles indicate significantly increased responses (q<0.05) from the baseline (Fz=0). Colored lines and shades represent mean responses ± S.E.M. across animals. Dashed lines mark time 0. **(i and j)** Peak Fz signal of MeA^Foxp2^ **(i)** and MeA^Dbx1^ cells **(j)** during the presentation of social and non-social stimuli. **(k)** Preference index (PI) of MeA^Foxp2^ and MeA^Dbx1^ cells to different social stimuli. For example, PI_male_ is calculated as (Fz_male_ − 0.5 × (Fz_female_ + Fz_pup_))/(Fz_male_ + 0.5×|Fz_female_ + Fz_pup_|). (d) Two-tailed unpaired t-test. (g-h) One sample t-test for each stimulus, corrected for multiple comparisons with a false discovery rate (FDR) 0.05. (i-j) One-way repeated-measures ANOVA followed by Tukey’s multiple comparisons test. (k) Two-way repeated measures ANOVA followed by Sidak’s multiple comparisons test. n = number of animals. Data are mean ± S.E.M.; *p<0.05, **p<0.01, ***p<0.001, ****p<0.0001.

We found that MeA^Foxp2^ cells in naïve male mice showed robust GCamp6 increases only during presentation of an adult male but not any other social stimuli (**Fig. 2e, g, i, k**). In contrast, MeA^Dbx1^ cells responded to all social stimuli with the highest activity increase during presentation of an adult female (**Fig. 2f, h, j-k**). Neither MeA^Foxp2^ nor MeA^Dbx1^ cells responded to a novel object, suggesting their social specific tuning (**Fig. 2e-f, i-j**). In addition to differential response selectivity, MeA^Foxp2^ and MeA^Dbx1^ cells also differed in their response dynamics. While MeA^Foxp2^cells responded slightly after the stimulus onset, defined as when the stimulus animal reached its minimum distance to the nose of the recording mouse, MeA^Dbx1^ cells significantly increased activity right at the onset of stimulus presentation (**Fig. 2g, h**). Furthermore, MeA^Foxp2^ cells returned to the baseline activity slowly (>10s) after removal of the male stimulus, while the MeA^Dbx1^ cell activity returned to the baseline quickly (< 3s) (**Figure 2g, h**). Overall, MeA^Foxp2^ cells showed male-specific and slow responses while MeA^Dbx1^ cells showed broad and fast responses to social cues (**Fig. 2g-k**). These results strongly support distinct response patterns of MeA^Foxp2^ and MeA^Dbx1^ cells to social stimuli independent of fighting or mating experience, with MeA^Foxp2^ cells displaying a select tuning to male cues.

### Distinct responses of MeA^Dbx1^ and MeA^Foxp2^ cells during social behaviors in freely moving male mice

Next, we examined responses of male MeA^Foxp2^ and MeA^Dbx1^ cells during social behaviors in freely moving male mice to address whether the cells increase activity only to sensory cues, e.g. during investigation, or also respond during the action phase of the behavior, e.g. attack and mount (**Extended Data Fig. 1a**). Prior to recording, all test animals went through repeated interactions with an adult male and a receptive female until they showed consistent aggression and sexual behaviors. During recording, an adult male intruder, a female, a pup and a novel object were introduced into the home cage of the recording mice, one at a time, with 5 minutes in between (**Extended Data Fig. 1b**). MeA^Foxp2^ cells showed significantly higher activity increase upon introduction of a male than any other social and non-social stimuli (**Fig. 3a-c, g, k**). During subsequent male investigation and attack, MeA^Foxp2^ cells also showed a significant activity increase (**Fig. 3h**). To address whether the attack response could largely be due to the continuous conspecific sensory input while attacking, instead of the attack per se, we separated investigation trials based on whether they were followed by attack or not. We found that activity increases during investigation-followed-by-attack trials was significantly higher than that during investigation-only trials at both investigation onset and offset (**Extended Data Fig. 1c)**. This result suggests that MeA^Foxp2^ respond during both sensory and action phases of aggression and the attack response is not simply due to temporally-linked sensory inputs. In contrast to the strong activity increase during male interaction, MeA^Foxp2^ cells showed either no change or slightly suppressed activity during female investigation and all phases of sexual behaviors (**Fig. 3b, h**). Similarly, no activity change was observed during pup interaction, supporting a highly male specific response of MeA^Foxp2^ cells (**Fig. 3c, h l**).

**Figure 3.**
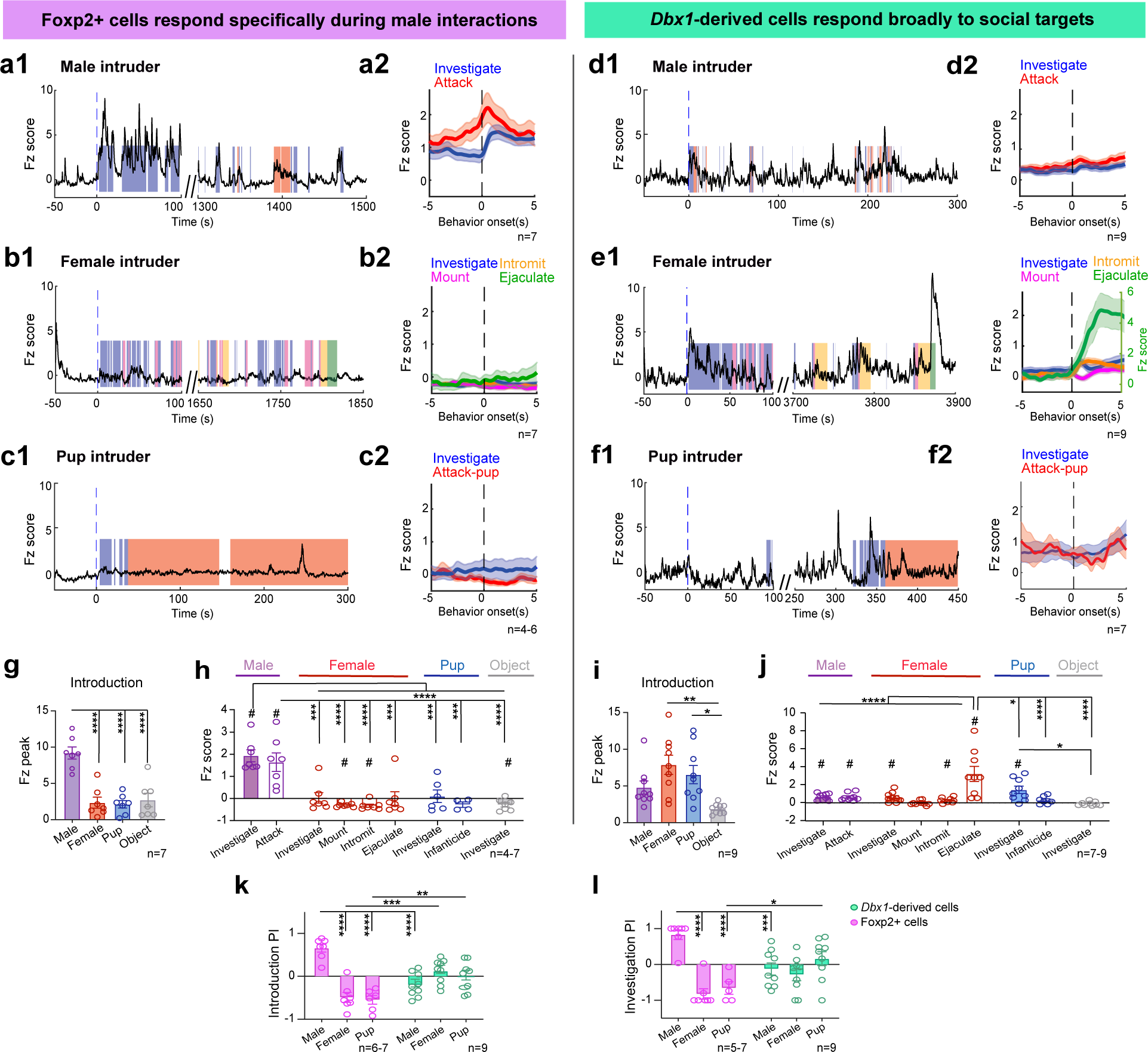
Differential response patterns of MeA^Foxp2^ and MeA^Dbx1^ cells during fighting and mating in socially experienced male mice. **(a-f)** Representative Ca^2+^ traces and peri-event histograms (PETHs) of MeA^Foxp2^ (**a-c**) and MeA^Dbx1^ cells (**d-f**) during interactions with male, female and pup stimuli. Dashed black lines in PETHs represent the behavior onset at time zero; blue lines in Ca^2+^ traces indicate time 0 when the intruder is introduced. **(g and i)** Introduction responses of MeA^Foxp2^ **(g)** and MeA^Dbx1^ cells **(i),** calculated as the peak Ca^2+^ signal within the first 100 sec after stimulus introduction. **(h and j)** Average Ca^2+^ responses of MeA^Foxp2^ **(h)** and MeA^Dbx1^ cells **(j)** during various behaviors towards various conspecific intruders and a novel object. **(k)** Preference indexes of MeA^Foxp2^ and MeA^Dbx1^ cells showing the relative introduction response magnitudes across different social stimuli. **(l)** Preference indexes of MeA^Foxp2^ and MeA^Dbx1^ cells denoting the relative investigation response magnitudes across different social stimuli. (g) One-way repeated-measures ANOVA followed by Tukey’s multiple comparisons test. (h, j) Mixed-effects analysis followed by Tukey’s multiple comparisons test. One sample t-test for each behavior, corrected for multiple comparisons with a false discovery rate (FDR) 0.05. (i) Friedman test followed by Dunn’s multiple comparisons test. (k-l) Mixed-effects analysis followed by Sidak’s multiple comparisons test. *n= 7* mice during male and female presentation for MeA^Foxp2^ group; *n=* 6 mice during pup investigation and *n=* 4 mice attacking pup for MeA^Foxp2^ group; *n=* 9 mice during male and female presentation for MeA^Dbx1^ group; *n=* 9 mice during pup investigation and *n=*7 attacking pup for MeA^Dbx1^ group. Data are mean ± S.E.M.; *p<0.05, **p<0.01, ***p<0.001, ****p<0.0001; # q<0.05.

In contrast to the response pattern of MeA^Foxp2^ cells, MeA^Dbx1^ cells in experienced male mice showed activity increases in response to all social stimuli (**Fig. 3d-f**). Upon initial intruder introduction, MeA^Dbx1^cells increased activity to all intruders with a slightly lower response to male intruder than females and pups (**Fig. 3i, k**). During investigation of a female, male and pup, MeA^Dbx1^cell activity increased to a similar extent (**Fig. 3j, l**). Although MeA^Dbx1^ cells also showed significant activity increase during inter-male attack, we did not find a difference in response between investigation-followed-by-attack trials and investigation-only trials, suggesting that MeA^Dbx1^ cell response during attack could be largely due to activity increases induced by sensory cues (**Fig. 3j, Extended Data Fig. 1d**). During copulation, the activity of MeA^Dbx1^ cells did not increase during mounting –a series of fast movements to establish an on-top position, but slightly increased during intromission (**Fig. 3e, j**). During ejaculation, MeA^Dbx1^ cells increased activity robustly, higher than the responses during any other behaviors (**Fig. 3e, j)**. No activity increase of MeA^Dbx1^ cells was observed when males attacked pups (**Fig. 3f, j**). Consistent with the response in head-fixed animals, neither MeA^Foxp2^ nor MeA^Dbx1^ responded during object investigation (**Extended Data Fig. 1e, f**), supporting the social-specific response patterns of the cells.

Overall, male MeA^Foxp2^ cells show highly specific responses during both the investigatory and action phases of behaviors towards a conspecific male whereas MeA^Dbx1^ cells appear to respond mainly to olfactory and possibly penile sensory inputs.

### Refinement of male MeA^Foxp2^ cell responses with adult social experience

In a subset of Foxp2^GCamP^ animals, we also performed Ca^2+^ recording during freely moving social interactions after head-fixed recording and before repeated social experience. Only one naïve male showed brief attack towards a male intruder and others only investigated the intruders. Consistent with recordings in head-fixed naïve animals, MeA^Foxp2^ cells responded specifically during male investigation (**Fig. 4a-f**). However, when we compared the response patterns of MeA^Foxp2^ cells in naïve vs. experienced animals, we noticed a clear difference. In comparison to naïve animals, activity of MeA^Foxp2^ cells in experienced animals increased faster and with higher reliability (**Fig. 4g-i**). In naïve animals, MeA^Foxp2^ cells responded (Z_increase_ > 1 during investigation) in approximately 40% of trials while this number increased to 60% in experienced animals (**Fig. 4j**). Among the responsive trials, the average latency to respond in experienced animals is approximately half of that in naïve animals (**Fig. 4k**). Overall, the mean activity increase during male investigation is significantly higher in experienced animals than in naïve animals although the male preference index (PI) did not differ between these two groups (**Fig. 4l-m**). The difference in response is not due to changes in investigatory behaviors: the average duration of investigation episodes was similar in naïve vs. experienced animals (**Fig. 4n**). It is worth noting that the difference in response patterns between naïve and experienced animals does not depend on the expression of aggression. Two experienced males did not attack the intruder during the recording (green circles in **Fig. 4j-n**) and their MeA^Foxp2^ cell responses were comparable to those in aggressive experienced males (**Fig. 4j-n**). These results suggest that although adult social experience is not required for the male specific responses of MeA^Foxp2^ cells, it refines the response by improving its consistency and temporal precision.

**Figure 4.**
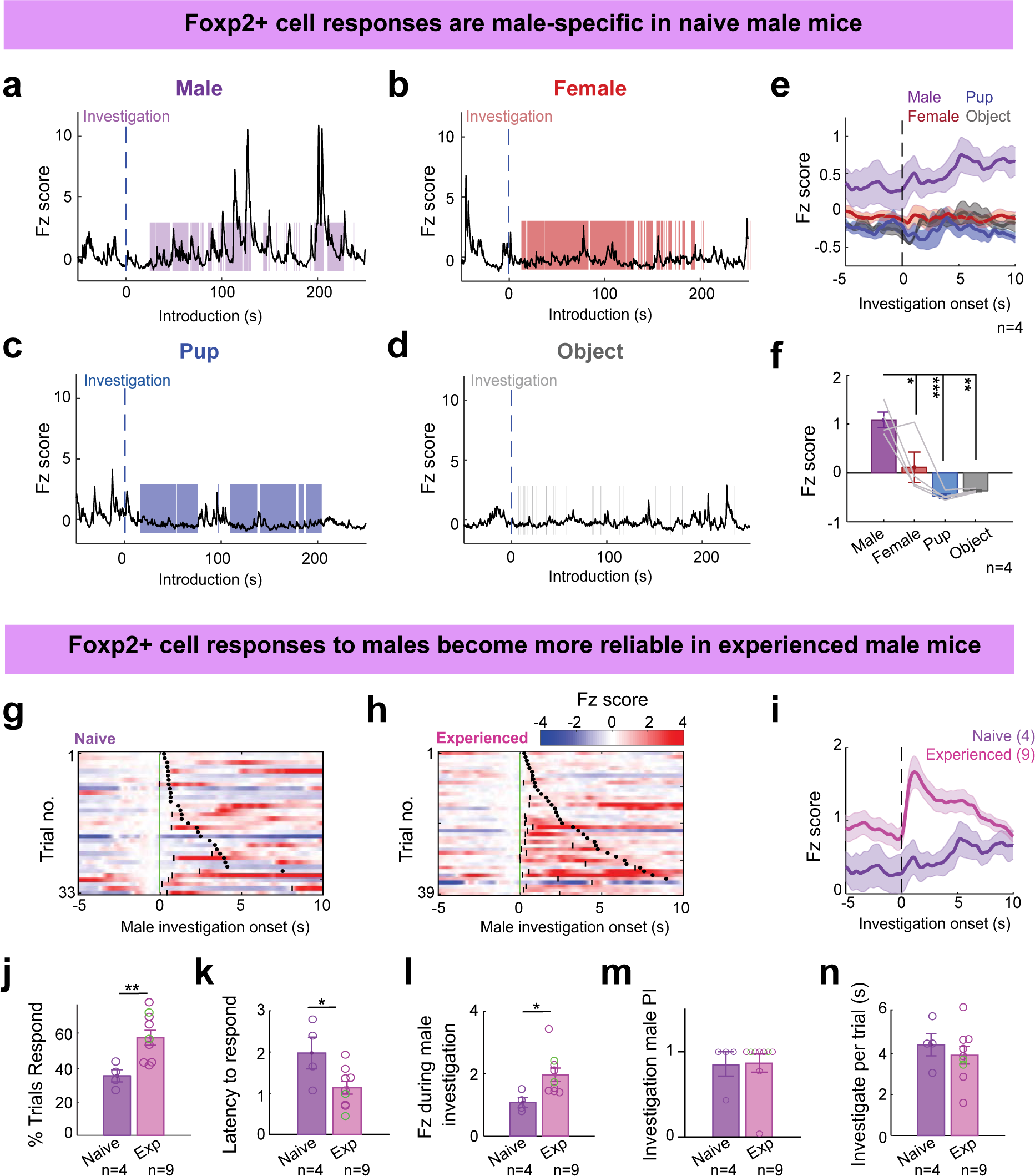
Comparison of MeA^Foxp2^ cell responses in naïve vs socially experienced male mice. **(a-d)** Representative Ca^2+^ traces of MeA^Foxp2^ cells during the presentation of a male **(a)**, female **(b),** pup **(c)** and object **(d)** in naïve male mice. **(e)** Average PETHs of MeA^Foxp2^ cell responses aligned to investigation onset in naïve male mice. The dashed black line represents the behavior onset at time zero. **(f)** Average Fz score of MeA^Foxp2^ cells during investigation of different stimuli in naïve male mice. **(g and h)** Representative heat-maps showing trial-by-trial Ca^2+^ signal (Fz-Fz at time 0) of MeA^Foxp2^ cells during investigation of a male intruder in naïve (**g**) and socially experienced **(h)** in a male mouse. Black short lines denote the time points when Fz >=1. Black dots denote the offsets of investigation. **(i)** Average PETHs of MeA^Foxp2^ cell responses aligned to investigation onset in naïve (purple) and socially experienced (pink) male mice. The dashed black line represents the behavior onset at time zero. **(j)** Percent of trials in which MeA^Foxp2^ cells show Fz>1 during male investigation in naïve and experienced male mice. **(k)** Latency of MeA^Foxp2^ cells to respond (Fz>1) in responsive trials. **(l)** Average Fz score of MeA^Foxp2^ cells during male investigation in naïve and experienced male mice. **(m)** Male preference index of MeA^Foxp2^ cell responses during investigation in naïve and experienced male mice. **(n)** Average male investigation duration in naïve and experienced male mice. Green circles in (j-n) represent male mice with repeated social experience but did not attack during the recording session. (f) One-way repeated-measures ANOVA followed by Tukey’s multiple comparisons test. (j-n) Two-tailed unpaired t-test. Parenthesis and n= number of mice. Data are mean ± S.E.M.; *p<0.05, **p<0.01, ***p<0.001.

### The male-specific response of male MeA^Foxp2^ cells exists before puberty

To further address whether the highly male specific MeA^Foxp2^ cells is developmentally hardwired or established through adult experience, we recorded the responses of MeA^Foxp2^ cells to social stimuli during early life. Puberty (P30-P38) is a critical development period when aggression starts to emerge^29-31^, thus we focused on MeA^Foxp2^cell responses before (P25), at the onset of (P30-32) and after puberty (P40-44). To achieve this goal, we injected Cre-dependent GCaMP6f virus into the MeA of P11 *Foxp2^cre^* mice and placed a 400-µm fiber just dorsal to the MeA at P24 (**Fig. 5a-b**). After a 24hr recovery window, we recorded the Ca^2+^ activity of MeA^Foxp2^ cells when the animals were exposed to an anaesthetized adult male or female mouse or a pup (**Fig. 5c**). To minimize the impact of social experience, all animals were singly housed post-weaning at P21. We found that in P25 juvenile male mice MeA^Foxp2^ cells already showed higher activity during interaction with an adult male than other social stimuli (**Fig. 5d**). At P30-32, a similar male-biased response was observed (**Fig. 5e**). At P40-44, the difference between male and female responses further increased and this trend continued at >P56 (**Fig. 5f**). Notably, the divergence of MeA^Foxp2^ cell responses to male and non-male cues over age appears to be mainly driven by a decrease in responses to females and pups (**Fig. 5h**). Behaviorally, juvenile (P25 and P30-32) and young adult (P40-44) males tended to investigate adult females more than adult males (**Fig. 5i**). However, this behavior difference did not explain the differential MeA^Foxp2^ responses to males and females as no correlation between response magnitude and time spent on investigation was found (**Fig. 5j**). Finally, the male preference index (PI) across all 4 time-points did not differ, supporting that the male-specific cell responses exist before puberty (**Fig. 5k**). Altogether, these results suggest that MeA^Foxp2^ cells are predisposed to preferentially respond to male-related sensory information even before puberty. We suggest that discriminability between male and non-male cues further improves after puberty by reducing responses to non-male cues.

**Figure 5.**
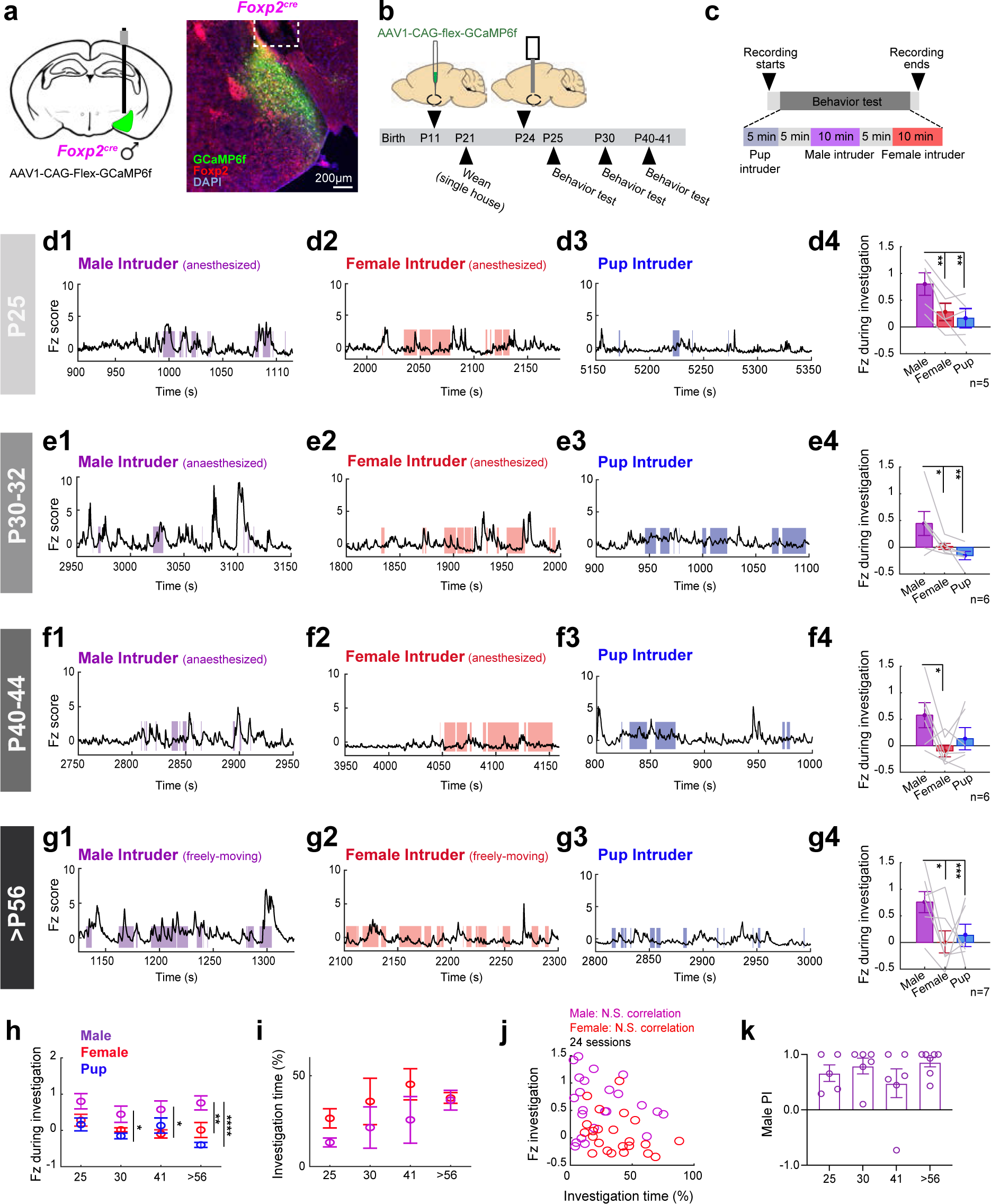
MeA^Foxp2^ cell responses before, during and after puberty in developing male mice. **(a)** Schematics of virus injection and a representative histology image indicating GCaMP6f expression (green), Foxp2 antibody (red) and DAPI (blue) staining in Foxp2^cre^ male mice. White dotted lines mark the fiber ending. **(b)** Timeline of virus injection, fiber placement and recordings. **(c)** Timeline of behavioral test during the recording day. Stimuli were presented in a pseudo-random fashion. **(d-g)** Representative Fz scored Ca^2+^ traces of MeA^Foxp2^ cells during interactions with an anesthetized (d1-f1) or freely-moving male (g1), an anesthetized (d2-f2) or freely-moving female (g2) or a pup (d3-g3) in a male mouse at different ages. Average Fz score during social investigation (d4-g4) of animals at different ages. **(h)** Average Fz score of MeA^Foxp2^ cell responses during male (purple), female (red) and pup (blue) investigation in mice at different ages. **(i)** Percent of time the test male spent investigating a male and female intruder. **(j)** No correlation between the Fz score during male or female investigation and the percent of time spent investigating in all recording sessions across ages. **(k)** Male investigation PI at different ages. (d4-g4) One-way repeated-measures ANOVA followed by Tukey’s multiple comparisons test. (h-i) Two-way repeated measures ANOVA followed by Sidak’s multiple comparisons test. (j) Pearson’s product-moment correlation coefficient (k) Kruskal-Wallis test. *n=*5 (P25), 6 (P30-32), 6 (P40-44) and 7 mice (>P56). Data are mean ± S.E.M.; *p<0.05, **p<0.01, ***p<0.001, ****p<0.0001.

### Differential inputs to MeA^Foxp2^ and MeA^Dbx1^ cells

Given the differential responses of MeA^Foxp2^and MeA^Dbx1^ cells to social cues, we next asked whether these the two populations receive different direct inputs. To test this, we used monosynaptic rabies virus tracing. We injected Cre-dependent or Flp-dependent AAVs expressing TVA-mCherry and rabies G protein into the MeA of *Foxp2^cre^*or *Dbx1^cre^;LSLFlpO* male mice, and four weeks later EnvA-ΔG rabies virus expressing GFP (**Fig. 6a-d)**. We found that the major inputs to MeA^Foxp2^ arise from other amygdala nuclei including posterior amygdala (PA), central amygdala (CeA) and BNST (**Fig. 6e-g**). In contrast, MeA^Dbx1^ cells receive inputs mainly from primary olfactory relays, including AOB, cortical amygdala (COA) and piriform cortex (Pir) (**Figure 6e, h, i**). Hypothalamus, mainly medial preoptic area (MPOA) and zona incerta (ZI), provided moderate inputs to both MeA^Foxp2^and MeA^Dbx1^ cells (**Fig. 6e-i**). Sparsely retrogradely labeled cells from both MeA^Foxp2^and MeA^Dbx1^ cells were also observed in hippocampus, striatum and pallidum (**Fig. 6e-i**).

**Figure 6.**
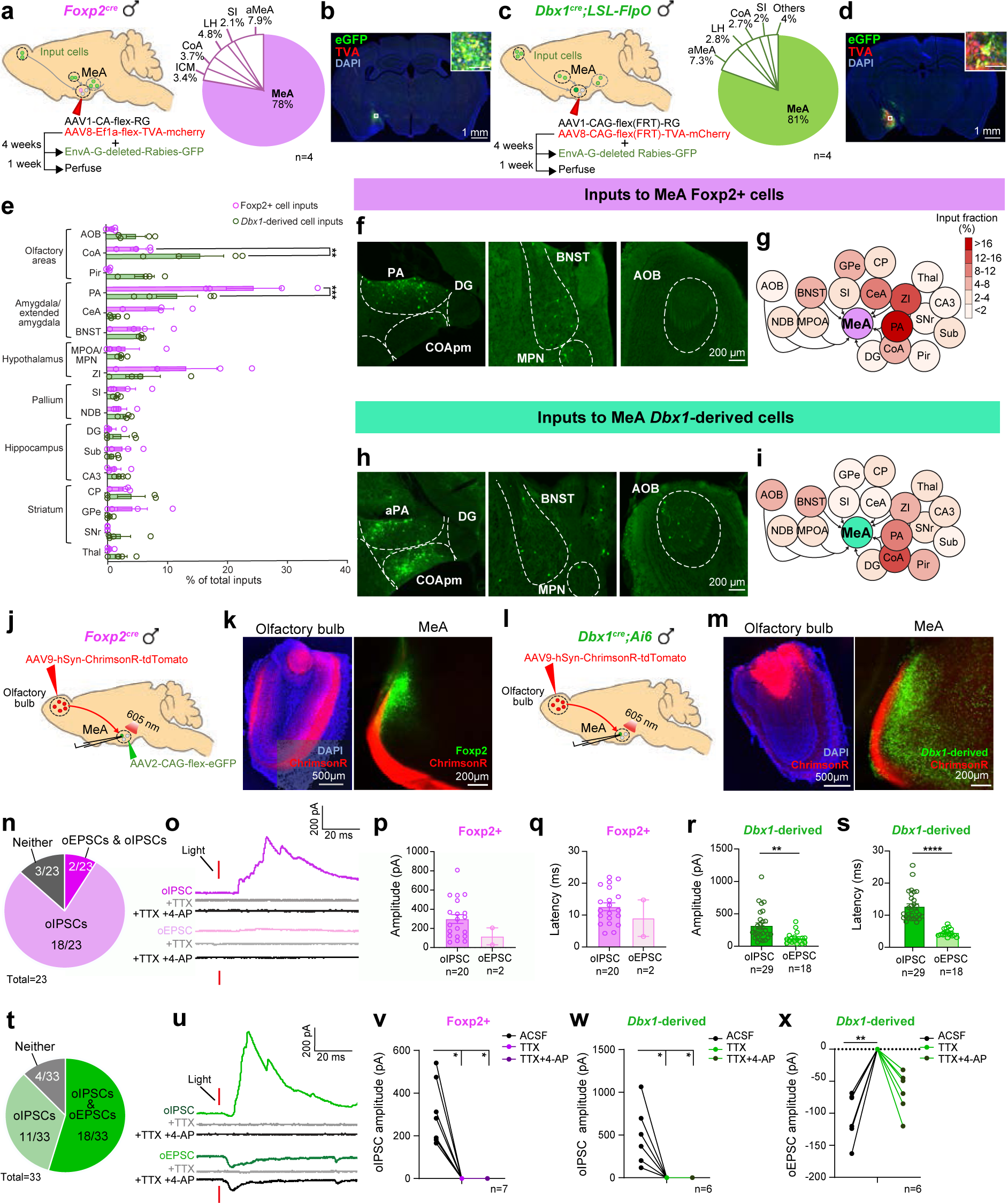
Differences in the anatomical and functional inputs of MeA^Foxp2^ and MeA^Dbx1^ cells for sensory processing. **(a)** Schematics showing timeline of monosynaptic retrograde rabies tracing of MeA^Foxp2^ cells. Pie chart showing the distribution of starter cells (mCherry+ eGFP+). **(b)** Representative image showing the location of starter MeA^Foxp2^ cells, denoting TVA-mCherry (red), Rabies-eGFP (green) and DAPI (blue) staining. Inset showing an enlarged view of boxed area. Scale bars: 1mm and 100µm (inset). **(c)** Schematics showing the retrograde monosynaptic tracing from MeA^Dbx1^ cells and the starter cell distribution. **(d)** Representative histology of the location of starter MeA^Dbx1^ cells in a Dbx1^cre^;LSL-FlpO mouse. Red: TVA-mCherry. Green: Rabies-eGFP, Blue: DAPI staining. Scale bars: 1mm and 100µm (inset). **(e)** Distribution of cells in various brain regions that are retrogradely labelled from MeA^Foxp2^ and MeA^Dbx1^ cells. **(f and h)** Representative histological images showing cells in various regions that are retrogradely labelled from MeA^Foxp2^ (f) or MeA^Dbx1^ (h) cells. **(g and i)** Overview of inputs into MeA^Foxp2^ (g) and MeA^Dbx1^ (i) cells. **(j and l)** Recording strategy to examine functional inputs from AOB to MeA^Foxp2^ (k) and MeA^Dbx1^ (l) cells. **(k and m)** Representative images showing ChrimsonR (red) expression in the olfactory bulb (OB) and ChrimsonR fibers in the MeA. Green: GFP expressed in Foxp2 (k) and Dbx1 (m) cells. Blue: DAPI staining. **(n and t)** Pie charts showing the distribution of synaptic responses of MeA^Foxp2^ (n) and MeA^Dbx1^ (t) cells to optogenetic activation of OB terminals. **(o and u)** Representative traces showing optogenetically (1 ms, 605 nm) evoked IPSCs (oIPCSs) and EPSCs (oEPSCs) before and after bath application of TTX and TTX + 4-AP. **(p-s)** Characterization of oIPSCs and oEPSCs in MeA^Foxp2^ and MeA^Dbx1^ cells, including amplitude **(p, r)** and latency **(q, s)**. **(v-w)** oIPSCs in both MeA^Foxp2^ (v) and MeA^Dbx1^ (w) cells were abolished by bath application of TTX and failed to recover after applying TTX+4-AP. **(x)** oEPSCs in MeA^Dbx1^ cells were abolished by TTX but recovered after TTX+4-AP application. SI: substantia innominate. NDB: diagonal band nucleus. DG: dentate gyrus. Sub: subiculum. CA3: field CA3. CP: caudoputamen. GPe: globus pallidus, external segment. SNr: substantia nigra, reticular part. Thal: thalamus. (e) Two-way ANOVA followed by Sidak’s multiple comparisons test; *n=*4 mice in each group. (r and s) Mann Whitney test. (v-x) Friedman test followed by Dunn’s multiple comparisons test. (n-s) n=23 cells from 3 male mice for MeA^Foxp2^ group; n=33 cells from 3 male mice for MeA^Dbx1^group; (v) n=7 cells from 3 male mice; (w, x) n=6 cells from 3 male mice. Data are mean ± S.E.M.; *p<0.05, **p<0.01, ***p<0.001, ****p<0.0001.

The lack of retrogradely labeled cells in AOB from MeA^Foxp2^ starter cells was particularly surprising given that the MeA is the primary target of the AOB, where the output neurons are excitatory (**Fig. 6e-g**)^4,32,33^. To further understand the inputs from the AOB to MeA^Foxp2^ cells, we performed optogenetic assisted circuit mapping from AOB to MeA^Foxp2^ and MeA^Dbx1^ cells. We expressed ChrimsonR-tdTomato in the olfactory bulb, virally labeled MeA^Foxp2^ cells with GFP (**Fig. 6j, k**) and visualized MeA^Dbx1^ cells using *Dbx1^cre^;Ai6* mice (**Fig. 6l, m**). 4-weeks after injection, we prepared brain slices containing the MeA and recorded the responses of GFP+ MeA^Foxp2^ and MeA^Dbx1^ cells to 605 nm 1-ms light pulses. Among a total of 23 MeA^Foxp2^ cells, we observed light evoked excitatory postsynaptic currents (oEPSCs) in only 2 cells while majority (18/23) of recorded cells showed light evoked inhibitory post-synaptic currents (oIPSCs) (**Fig. 6n, o**). In contrast, 18/33 MeA^Dbx1^ cells showed oEPSCs and the vast majority (29/33) showed oIPSCs (**Fig. 6t, u**). The oIPSCs of MeA^Dbx1^ and MeA^Foxp2^ cells were similar in magnitude and both were of long latencies (>10ms) (**Fig. 6p-s**). Bath application of TTX or TTX+4-AP completely abolished oIPSCS in both populations, suggesting that both MeA^Foxp2^and MeA^Dbx1^ cells receive polysynaptic inhibitory inputs (**Fig. 6o, u, w**). oEPSCs of MeA^Dbx1^cells are of shorter latency (∼4 ms) than oIPSCs (**Fig. 6s**) and bath application of TTX+4-AP did not abolish oEPSCs, supporting that AOB cells provide monosynaptic excitatory inputs to MeA^Dbx1^ cells (**Fig. 6u, x**).

These results confirmed that AOB targets MEA^Foxp2^ and MeA^Dbx1^ cells differently, consistent with the idea that the distinct *in vivo* responses of these two populations are hardwired. The fact that MeA^Foxp2^ cells receive minimum direct inputs from the AOB and other primary olfactory relays suggests that sensory information reaching MeA^Foxp2^ cells could be more processed, which may explain the higher response selectivity of MeA^Foxp2^ cells than MeA^Dbx1^ cells.

### MEA^Foxp2^ cells are sufficient to promote inter-male aggression in naïve mice

To understand the functional importance of MeA^Foxp2^ and MeA^Dbx1^ cells in social behaviors, we bilaterally injected Cre- and Flp-dependent hM3Dq viruses into the MeA of *Foxp2^cre^* and *Dbx1^cre^;LSL-FlpO* naïve male mice respectively (Foxp2^hM3Dq^ and Dbx1^hM3Dq^) (**Fig. 7a, b**). Control animals were injected with mCherry virus in the MeA (Foxp2^mCherry^ and Dbx1^mCherry^). Three weeks later, we intraperitoneally (i.p.) injected saline and clozapine-N-oxide (CNO) on two separate days and 30 mins later introduced a pup, an adult male and a female intruder into the cage sequentially, each for 5 to 10 minutes, with 5 minutes in between (**Fig. 7c**). While only 4/10 Foxp2^hM3Dq^ male mice attacked a male intruder after saline injection, all Foxp2^hM3Dq^ males attacked the intruder repeatedly after CNO injection (**Fig. 7e**). In comparison, only 4/8 Foxp2^mCherry^ initiated attack after CNO injection (**Fig. 7e**). The total attack duration of Foxp2^hM3Dq^ males significantly increased after CNO injection (**Fig. 7f)** although the latency to attack did not decrease in animals that attacked on both days (**Extended Data Fig. 2a**). Possibly due to increased aggression, Foxp2^hM3Dq^ mice spent less time investigating the male intruder after CNO injection (**Fig. 7g**). No changes in locomotion were observed in Foxp2^hM3Dq^ males after CNO injection suggesting that increases in attack was not due to an increase in general arousal (**Extended Data Fig. 2b**). Additionally, the increased aggression is adult male-specific as we did not observe an increase in infanticidal behavior after activating MEA^Foxp2^ cells (**Extended Data Fig. 2c)**. The overall pup interaction was also unchanged (**Extended Data Fig. 2d)**. Similarly, male sexual behaviors, including female investigation, mounting and intromission, were not affected by MEA^Foxp2^ activation (**Extended Data Fig. 2e-k**). Control Foxp2^mCherry^ animals showed no significant change in any social behavior after CNO injection in comparison to saline injection (**Fig. 7d-g, Extended Data Fig. 2a-k**).

**Figure 7.**
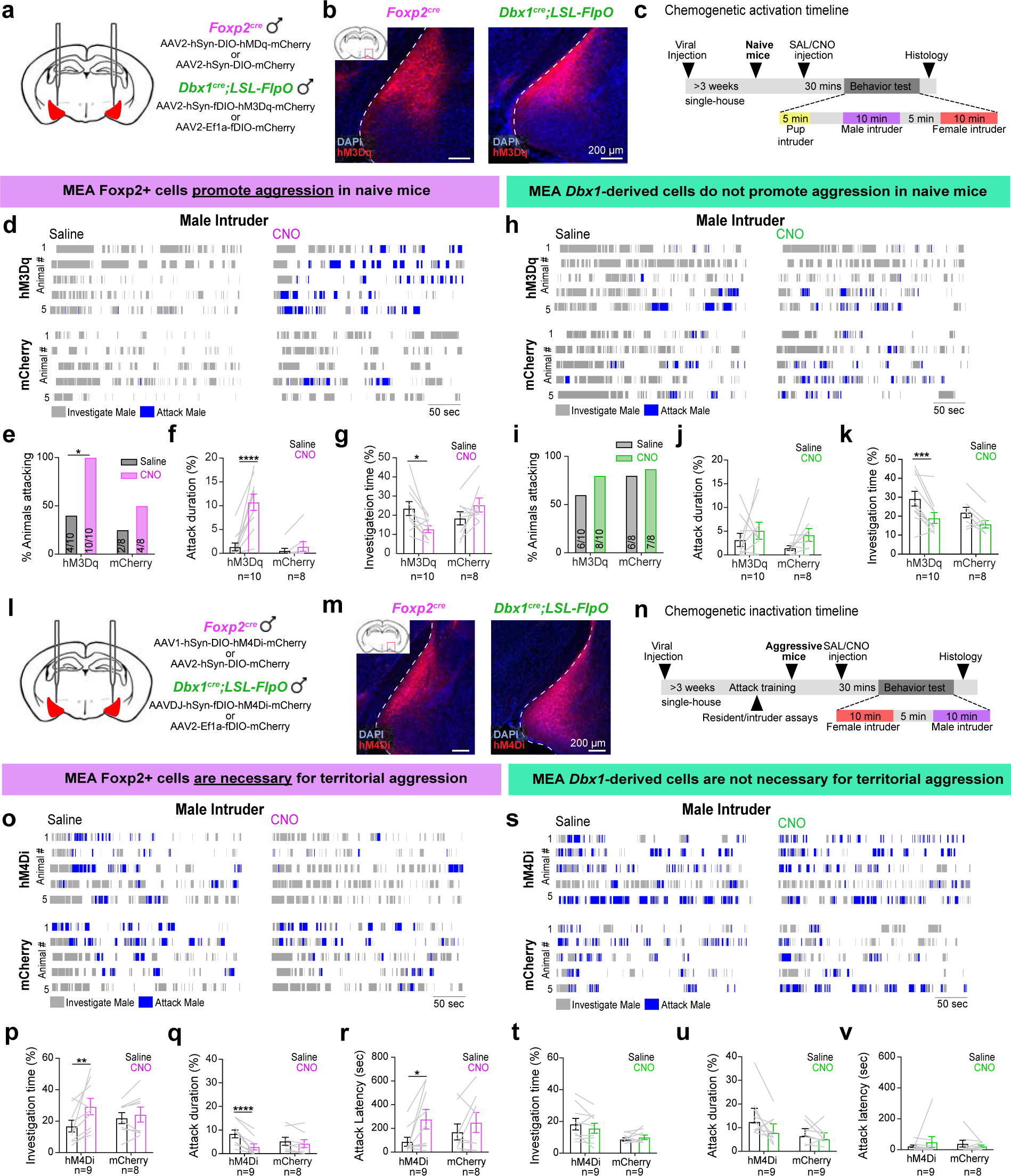
MeA^Foxp2^ cells are necessary and sufficient for territorial aggression, while MeA^Dbx1^ cells are not. **(a)** Strategies for chemogenetic activation of MeA^Foxp2^ and MeA^Dbx1^ cells. **(b)** Representative histological images of hM3Dq (red) expression in the MeA of Foxp2^cre^ and Dbx1^cre^;LSL-FlpO mice. Blue: DAPI. **(c)** Experimental timeline of chemogenetic activation experiments. **(d)** Representative raster plots showing behaviors towards male intruders of 5 Foxp2^hM3Dq^ and 5 Foxp2^mCherry^ male mice after i.p. injection of saline or CNO. **(e)** Percentage of Foxp2^hM3Dq^ and Foxp2^mCherry^ male mice that attacked a male intruder after saline or CNO injection. **(f-g)** Percent of time Foxp2^hM3Dq^ and Foxp2^mCherry^ mice spent attacking (f) and investigating (g) a male intruder. **(h-k)** Follow conventions in d-g. CNO injection into Dbx1^hM3Dq^ mice caused a reduction in social investigation but did not change aggressive behaviors towards a male intruder. **(l)** Strategies for chemogenetic inactivation of MeA^Foxp2^ and MeA^Dbx1^ cells. **(m)** Representative histological images showing hM4Di (red) expression in the MeA of Foxp2^cre^ and Dbx1^cre^;LSL-FlpO mice. Blue: DAPI. **(n)** Experimental timeline of chemogenetic inactivation experiments. **(o)** Representative raster plots showing the behaviors of 5 Foxp2^hM4Di^ and 5 Foxp2^mCherry^ mice after i.p. injection of saline or CNO in the presence of a male intruder. **(p-r)** Percent of time Foxp2^hM4Di^ and Foxp2^mCherry^ male mice spent investigating (p) and attacking (q) a male intruder, and the latency to first attack (r). **(s-v)** Follows the conventions in o-r. CNO injection into Dbx1^hM4Di^ or Dbx1^mCherry^ mice did not change any male-directed behaviors in comparison to those after saline injection. (e, i) McNemar’s test. (f, g, j, k, p-r, t-v) Two-way repeated measures ANOVA followed by Sidak’s multiple comparisons test. n = number of animals. Data are mean ± S.E.M.; *p<0.05, **p<0.01, ***p<0.001, ****p<0.0001.

We found that Dbx1^cre^;LSL-FlpO male mice tend to be more aggressive than Foxp2^cre^ male mice possibly due to their slight difference in genetic background^27,28^. Specifically, majority of Dbx1^hM3Dq^ and Dbx1^mCherry^ animals attacked the intruder during the first encounter (after saline injection) and nearly all animals attacked the intruder during the second encounter (after CNO injection) (**Fig. 7h, i**). Importantly, there is no difference between Dbx1^hM3Dq^ and Dbx1^mCherry^ groups in the percentage of animals that attacked (**Fig. 7i**). The latency to attack and attack duration also did not differ on CNO- and saline-injected days in both Dbx1^hM3Dq^ and Dbx1^mCherry^ groups (**Fig. 7j, Extended Data Fig. 3a**) although Dbx1^hM3Dq^ male mice investigated the male intruder less after CNO injection (**Fig. 7k**). Activating MeA^Dbx1^ cells didn’t change the probability of infanticide, male sexual behaviors or locomotion significantly (**Extended Data Fig. 3b-k**). Thus, MEA^Foxp2^ cells can specifically drive inter-male aggression in even non-aggressive naïve male mice whereas activating MEA^Dbx1^ cells does not promote any specific social behaviors to a significant level.

### MeA^Foxp2^ cells but not MeA^Dbx1^ cells are necessary for inter-male aggression in experienced animals

We next asked whether MeA^Foxp2^and MeA^Dbx1^ cells are necessary for social behaviors, including inter-male aggression. We injected Cre- and Flp-dependent hM4Di-mCherry into the MeA of *Foxp2^cre^* and *Dbx1^cre^;LSLFlp* male mice respectively (Foxp2^hM4Di^ and Dbx1^hM4Di^). Control animals were injected with mCherry virus (**Fig. 7l, m**). Three weeks after viral injection, all animals went through repeated resident-intruder test until they showed stable level of aggression (**Fig. 7n**). Then, we i.p. injected saline and CNO on separate days in a randomized order and 30 minutes later tested the behaviors against a male and then a receptive female intruder, each for 10 minutes (**Fig. 7n**). After CNO injection, Foxp2^hM4Di^ mice spent more time investigating the male intruders and less time attacking the intruder (**Fig. 7o-q**). The latency to first attack increased significantly (**Fig. 7r**). Foxp2^mCherry^ mice showed no difference in male investigation or attack duration between CNO and saline injected days (**Fig. 7o-r**). In contrast, CNO injection in Dbx1^hM4Di^ mice did not result in significant changes in male investigation, aggressive behaviors, or latency to attack (**Fig. 7s-v**). CNO injection in Foxp2^hM4Di^ or Dbx1^hM4Di^ mice caused no change in female investigation or any aspects of male sexual behaviors except an increase in mount number in both Dbx1^hM4Di^ and Dbx1^mcherry^ groups (**Extended Data Fig. 2l-r, 3l-r**). These results suggest that MeA^Foxp2^ cells are required specifically for inter-male aggression while MeA^Dbx1^ cells are not.

### Differential outputs of MeA^Dbx1^ and MeA^Foxp2^ cells

As MeA^Dbx1^ and MeA^Foxp2^ cells play differential roles in driving social behaviors, presumably through their differential impact on downstream cells, we next asked whether these two MeA subpopulations differ in their projections using anterograde virus tracing (**Fig. 8a-d**). We observed that both MeA subpopulations project mainly to other extended amygdala areas, such as PA, COA and posterior BNST (BNSTp), and medial hypothalamus (MH) (**Fig. 8e-i, Extended Data Fig. 4**). Although the average density of projections originating from MeA^Dbx1^ and MeA^Foxp2^ did not differ in any brain region (**Fig. 8e**), we observed that MeA^Dbx1^ and MeA^Foxp2^ showed differential projection patterns in the pBNST and MH. While MeA^Dbx1^ cells targeted primarily the principal nucleus of the BNST (BNSTpr), MeA^Foxp2^ cells projected to both principal and interfascicular parts of the BNST (BNSTpr and BNSTif) (**Fig. 8j, k**). In the MH, we observed that MeA^Dbx1^ cells generally provided more inputs to structures in the anterior MH (Bregma level: 0.14– −0.75 mm) than posterior MH (Bregma level: −1.25-−2.15 mm) whereas MeA^Foxp2^ cells projected to the anterior and posterior MH similarly (**Fig. 8l, m**).

**Figure 8.**
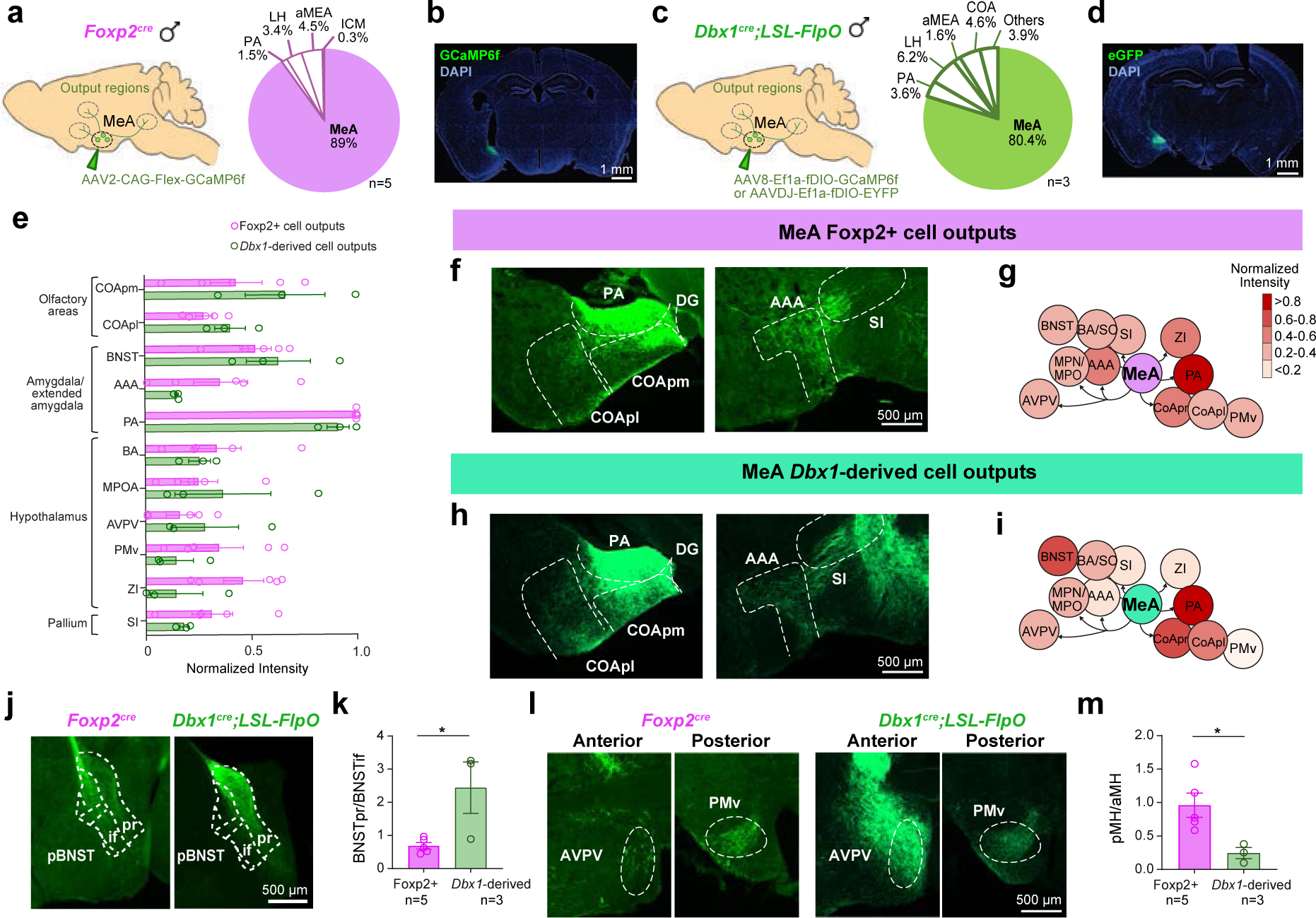
Outputs of MeA^Foxp2^ and MeA^Dbx1^ cells. **(a and c)** Strategies for anterograde viral tracing of MeA^Foxp2^ (a) and MeA^Dbx1^ (c) cells. Pie charts showing the distribution of primary infected cells. **(b and d)** Representative histological images showing the primary infected cells in Foxp2^cre^ (b) and Dbx1^cre^;LSL-FlpO mice (d).Green: eGFP or GCaMP6f expression. Blue: DAPI staining. **(e)** The intensity of MeA^Foxp2^ and MeA^Dbx1^ projection field in various regions, calculated as the average pixel intensity in a given region divided by the maximum average value across all regions. **(f and h)** Representative histological images showing MeA^Foxp2^ (f) and MeA^Dbx1^ (h) projections at various downstream regions. **(g and i)** Overviews of MeA^Foxp2^ (g) and MeA^Dbx1^ (i) cell outputs. **(j)** Images showing MeA^Foxp2^ and MeA^Dbx1^ cell outputs at pBNST. **(k)** The intensity of fibers, originating from MeA^Foxp2^ and MeA^Dbx1^ cells, at BNSTpr over that in BNSTif. **(l)** Representative histological images showing MeA^Foxp2^ and MeA^Dbx1^ projections at the anterior and posterior medial hypothalamus (aMH and pMH). **(m)** The intensity of fibers, originating from MeA^Foxp2^ and MeA^Dbx1^ cells, at pMH over that in aMH. pMH: Bregma −1.25 mm to −2.15mm; aMH: Bregma 0.14mm to −0.75mm. COApm: posteromedial cortical amygdala; COApl: posterolateral cortical amygdala; AAA: anterior amygdalar area; BA: bed nucleus of the accessory olfactory tract; AVPV: anteroventral periventricular nucleus; PMv: ventral premammillary nucleus. (e) Two-way repeated measures ANOVA followed by Sidak’s multiple comparisons test. (k, m) Two-tailed unpaired t-test. n = number of mice. Data are mean ± S.E.M.; *p<0.05.

## Discussion

In this study, we showed that two MeA subpopulations with different development lineages play distinct roles in social behaviors. They receive differential anatomical inputs and are responsive to distinct conspecific sensory cues. The male specific responses of MeA^Foxp2^ cells exist prior to puberty and aggression onset, suggesting that it is largely developmentally hardwired. The reliability, but not specificity, of MeA^Foxp2^ cell responses improve with adult social experience, demonstrating distinct roles of nature vs. nurture in establishing the social behavior circuit.

### MeA^Foxp2^ and MeA^Dbx1^ cell activity and function in innate social behaviors

Our previous work had identified two developmentally distinct GABAergic MeA subpopulations, marked by the expression of Dbx1 and Foxp2^9,24^. These two subpopulations differ in their sex steroid hormone receptor expression, ion channel composition, and intrinsic electrophysiological properties^9,34^. Our current study further revealed their distinct functions in social behaviors that are well matched with their connectivity and *in vivo* response patterns. These results suggest that social circuits at the MeA could be largely hardwired according to transcription factor-defined genetic programs.

MeA^Foxp2^ cells responded strongly during both male investigation and attack. Importantly, MeA^Foxp2^ show higher responses during male investigation when it is followed by attack, suggesting that the attack response is not simply due to sensory inputs when the animals are in close proximity. These results suggest that MeA is not merely a sensory relay, instead, it could serve a direct role in driving consummatory social actions. Consistent with this hypothesis, activation of MeA^Foxp2^ cells promoted male-directed attack even in inexperienced non-aggressive male mice.

Hong et. al. showed originally that optogenetic activation of MeA GABAergic cells can induce attack^16^ but a recent study found the manipulation was ineffective^35^. These opposite results appear to be caused by the different photocurrent magnitude of the chosen opsin: only ChR2 variants with large, but not small photocurrents, can induce attack from MeA GABAergic cells^16^ ^35,36^. Given that MeA^Foxp2^ cells have lower resting membrane potential, lower input resistance and lower spontaneous firing rate in comparison to MeA^Dbx1^ cells^9^, we speculate that MeA^Foxp2^ cells could be relatively hard to activate, which may explain why strong optogenetic activation is needed to induce attack from the MeA. Importantly, as activating MeA^Dbx1^ cells, which are three times more abundant than MeA^Foxp2^ cells, do not elicit attack, our study clearly argues that aggression generation requires activation of specific, instead of a random subset of MeA GABAergic cells.

In contrast to MeA^Foxp2^ cells, MeA^Dbx1^ cells are tuned to broad social cues, including those from males, females and pups, but respond minimally during the action phase of social behaviors. They are suppressed during mounting and showed similar responses in male investigation-only and investigation-followed-by attack trials. Consistent with their lack of activity change during social actions, inactivation of MeA^Dbx1^ cells does not impair male sexual and aggressive behaviors. Given the response pattern of MeA^Dbx1^ cells, we consider their main role as to process social cues during the investigatory phase. However, animals with inactivated MeA^Dbx1^ cells properly directed their attack towards males and mount towards females, suggesting that MeA^Dbx1^ cells are dispensable for sex discrimination. The lack of behavior deficits after MeA^Dbx1^ manipulation is possibly due to the existence of other extended amygdala populations that can readily distinguish male and female cues during social investigation, e.g. MeA^Foxp2^ and aromatase cells in BNSTpr^37^.

Previous work has shown that MeA GABAergic cells are activated during pup-directed attack and can promote infanticide^22^. However, neither MeA^Foxp2^ nor MeA^Dbx1^ cells increased activity during pup-directed aggression or affected infanticide when being artificially activated. This result suggests that MeA^Foxp2^ is specialized for aggression towards males. Other GABAergic subclasses likely exist for driving infanticide and remain to be identified.

### Developmentally wired vs. experientially wired

There is an ongoing debate whether the responses of cells in the SBN are developmentally hardwired or established through adult social experience. In the VMHvl, an essential region for male aggression^38-40^, individual cell responses to male and female cues overlap extensively in naïve adult male mice and only diverge after repeated interaction with females^41^. In contrast, aromatase expressing cells in male BNSTpr were found to preferentially respond to female cues over male cues even in naïve animals^37^. Ca^2+^ imaging in the MeA revealed that approximately half of MeA cells are tuned to one stimulus in naïve animals and after sexual experience the proportion of cells that are responsive to the opposite sex increases, denoting experience-dependent activity refinement^10^. In our study, MeA^Foxp2^ cells showed strong male-biased responses in naïve animals suggesting that male olfactory inputs are developmentally wired to target MeA^Foxp2^ cells. However, the responses of MeA^Foxp2^ cells in naïve males are slow and unreliable and only become fast and consistent after repeated social interactions, suggesting that adult social experience plays an important role in refining the hardwired circuit to improve its input (sensory cue)-output (spiking) transformation efficiency.

How is the male specific response of MeA^Foxp2^ cells achieved during development? The classical ‘organization/activation’ model states that gonadal hormones act in two phases to establish sex-specific circuits^42-44^. First, during the organization stage, gonadal hormones during prenatal development set up the basic structure and connection of the circuit. Then, the circuits are activated by gonadal hormones during puberty to generate appropriate sex-specific social behaviors. In male mice, puberty occurs between P30 and P40 when testosterone spikes and aggression emerges^29,42^. Previous single-unit recordings found that social response selectivity of MeA cells in anaesthetized juveniles (P18-21) is lower than that in adults, suggesting that sex-hormone mediated circuit “activation” during puberty is important for establishing adult MeA responses^6^. Here, our longitudinal recording revealed male-biased responses of MeA^Foxp2^ cells even before puberty, suggesting that the male cues have already been wired preferentially to MeA^Foxp2^ cells during the organization stage. After puberty, MeA^Foxp2^ cells show enhanced male-biased responses due to decreased responses to other social cues, e.g. female. As MeA^Foxp2^ cells do not express aromatase, which is important for the activation of the male territorial aggression circuit during development, and only express low levels of steroid hormone receptors^9^, hormone actions onto MeA^Foxp2^ cells might be limited. Therefore, we speculate that changes in the synaptic inputs that suppress non-male related inputs could be the main mechanism responsible for the increased specificity after puberty. As the animals become full adults and acquire social experiences, the response to non-male cues remains low while responses to males continue to increase. Altogether, we propose that the response specificity of MeA^Foxp2^ cells during development is achieved through a multistage process, including pre-pubertal hardwiring, pubertal refinement, and adult social experience-dependent potentiation. Future microcircuit studies could help further validate this model and its potential generality in the SBN.

### Social behavior circuits beyond MeA

In mice, olfactory inputs are the most essential for determining the identity of a conspecific, e.g. its sex, age, social ranking and health state (e.g. sickness)^45^. Since MeA^Foxp2^ cells receive little direct input from the AOB and other primary olfactory relays, we speculate that MeA^Foxp2^ cells obtain highly “processed” olfactory information from the PA. Recent work from our group and others revealed that PA cells that project to the VMHvl are crucial for territorial aggression and these cells are activated during both male investigation and attack^46,47^. The PA also projects strongly to MeA; however, whether this projection is essential for aggression remains to be explored. On the contrary, MeA^Dbx1^ cells receive abundant inputs from AOB and other primary olfactory processing regions, which could be responsible for the broad and fast responses of MeA^Dbx1^ cells to various social cues.

At the output level, MeA^Dbx1^ and MeA^Foxp2^ cells project to distinct pBNST subnuclei: MeA^Dbx1^ cells project primarily to the BNSTpr while MeA^Foxp2^ cells project mainly to the BNSTif. Miller et al recently demonstrated that MeA cells that express D1R primarily targets the BNSTif and activating MeA^D1R^-BNST projections increased territorial aggression towards a conspecific^48^. This highlights the relevant role of BNSTif in aggression, and a potential downstream mechanism by which MeA^Foxp2^ cells mediate aggressive action. Additionally, MeA^Dbx1^ cells project mainly to anterior MH while MeA^Foxp2^ project similarly to anterior and posterior MH. Given that anterior MH, such as the anteroventral periventricular nucleus (AVPV) and the MPN, is most relevant for sexual behaviors, while the posterior MH, such as the VMHvl and the ventral premammillary nucleus (PMv), is central for male aggression^39,40,49^, the stronger projection of MeA^Foxp2^ cells to posterior MH in comparison to MeA^Dbx1^ is consistent with the essential role of MeA^Foxp2^ cells in male aggression.

### Transcription factor code in the limbic system

Analogous to the transcriptional code observed in the spinal cord and basal ganglia for cellular specificity of intrinsic physiology, connectivity and motor control, we suggest a transcription factor code in the limbic system by which distinct sets of transcriptionally-defined subpopulations differing in their intrinsic properties and connectivity mediate diverse behavioral functions^50,51^. Previous work investigating the LIM-homeodomain family of transcription factors, found two distinct MeA subpopulations expressing Lhx6 and Lhx9 that are relevant for reproduction and predator defense behaviors respectively^7^. Our results provide an *in vivo* understanding of additional transcriptionally defined subpopulations relevant for specific social behaviors, such as aggression.

A role of MeA^Foxp2^ in generating aggression is consistent with a role of Foxp2 in the basal ganglia, cerebellum and cortex in modulating motor actions^52^. Previous work has shown that Foxp2 expression is required for distinct components of motor actions: *Foxp2* in the cerebellum is essential for appropriate response-time and motor execution, in the striatum it decreases response variability, while in the cortex it is needed for appropriate motor performance^52^. In addition, given its expression in sensory processing regions, such as thalamus and association cortex, Foxp2 has been considered essential for sensorimotor integration of external cues, particularly auditory, for appropriate limbic movements^53-55^. Similar to cortical and subcortical regions, MeA^Foxp2^ cells are relevant for attack, a stereotyped aggressive action. Importantly, here we demonstrate that MeA^Foxp2^ involvement in motor generation appears to be tightly linked to its direct role in processing male specific olfactory inputs, going beyond its known role in auditory information processing. In parallel to our findings regarding MeA^Foxp2^ and MeA^Dbx1^, the globus pallidus externa (GPe) has been shown to comprise different inhibitory subpopulations arising from distinct progenitor pools in the medial and lateral/central ganglionic eminences, including an arkypallial subtype that expresses Foxp2 (GPe^Foxp2^) with intrinsic biophysical properties similar to that of MeA^Foxp2^ cells, and a prototypical subtype with intrinsic properties similar to those of MeA^Dbx1^ cells^9,34,51^. Nevertheless, differences across regions remain. The GPe^Foxp2^ functions primarily as a movement generator, similar to cerebellum and cortex, while MeA^Foxp2^ cells do not encode moment-to-moment movement, but instead, a specific behavior output (i.e. attack) comprised of a complex sequence of actions.

Overall, our study identified a developmentally hardwired circuit at the MeA that transforms male conspecific cues to attack command. It revealed the distinct contribution of development vs. experience in social information processing and highlighted a lineage-based organization strategy that enables the same SBN to drive diverse social behaviors^2^.

## Methods

### Mice

All animal procedures were approved by the Institutional Animal Care and Use Committee (IACUC) of NYU Langone Health. Adult experimental and stimulus mice were housed under a 12 hr light-dark cycle (10a.m. to 10p.m. dark) with water and food *ad libitum*. After surgical procedures, all experimental animals were single-housed. The *Foxp2^cre^* mice were originally provided by Dr. Richard Palmiter (now Jackson stock no. 030541)^27^. The *Dbx1^cre^* mice were originally provided by Dr. Alessandra Pierani and crossed to the Flp excised and Cre-inducible *LSL-FlpO* mouse line or to the Ai6 mouse line (Jackson stock no. 028584 and no. 007906 respectively)^25,26,28^. Both *Foxp2^cre^* and *Dbx1^cre^* mice are black, while the fur color of *LSL-FlpO* mice is agouti. Stimulus animals were C57BL/6N and 129S4/SvJae group-housed females, pups (P1-P7) and group-housed BALB/c males purchased from Charles River and bred in-house. Females were considered receptive if an experienced male was able to mount and intromit the female in at least 3 attempts.

### Viruses and stereotaxis surgery

For fiber photometry experiments, we injected 100nl of AAV2-CAG-Flex-GCaMP6f (2.21 × 10^13^ vg/ml or 1.82×10^12^ vg/ml; UPenn viral core) unilaterally into the MeA (AP: −1.5mm, ML= 2.15mm, DV: −5.1mm) of Foxp2^cre+/-^ male mice. For Dbx1^cre+/-^;FlpO^+/-^ mice we injected either 100nl AAV8-Ef1a-fDIO-GCaMP6f (1×10^13^ vg/ml; kindly provided by Dr. Uchida) or 120nl of mixed AAV9-Ef1a-fDIO-Cre (2.5×10^13^ vg/ml; Addgene) and AAV2-CAG-Flex-GCaMP6f (1:2; 2.21×10^13^ vg/ml; UPenn viral core) or 150nl of AAV2-Ef1a-fDIO-GCaMP6f (4.1 × 10^12^ vg/ml;UNC vector core) into the MeA. For fiber photometry recordings in Foxp2^cre+/-^ juvenile mice we injected 100nl of AAV1-CAG-Flex-GCaMP6f (9.4 × 10^12^ vg/ml; UPenn viral core) unilaterally into the developing MeA (AP: −0.7mm, ML= 2.03mm, DV: −5.05mm). For chemogenetic experiments, we bilaterally injected either 400-600nl of AAV1-Ef1a-DIO-hM4D(Gi)-mcherry, 150nl of AAV2-hSyn-DIO-hM3D(Gq)-mcherry or 150-600nl of AAV2-hSyn-DIO-mCherry (3×10^12^ vg/ml, 5.1×10^12^ vg/ml and 5.6×10^12^ vg/ml, respectively; Addgene and UNC Vector Core) into the MeA of Foxp2^cre+/-^ mice. For chemogenetic experiments in Dbx1^cre+/^-;FlpO^+/-^ mice, we injected 300nl AAVDJ-hSyn-fDIO-hM4D(Gi)-mCherry, 50-60nl AAV2-Ef1a-fDIO-hM3D(Gq)-mCherry (Vigene) and 60-120nl AAV2-Ef1a-fDIO-mCherry (2.65×10^13^ vg/ml, 1.84×10^13^ vg/ml and 1.1×10^13^ vg/ml, respectively; Addgene). For monosynaptic retrograde rabies experiments in Foxp2^cre+/-^ mice we injected unilaterally into the MeA 250-500nl of mixed AAV1-CA-Flex-RG and AAV8-Ef1-Flex-TVA-mCherry (1:1; 3×10^12^ vg/ml and 5.4 × 10^12^ vg/ml; UNC vector core) and 4 weeks later 800nl EnvA G-Deleted Rabies-eGFP (Salk viral vector core). For monosynaptic retrograde rabies experiments in Dbx1^cre+/^-;FlpO^+/-^ mice we injected mixed 110-120nl AAV8-Flex(FRT)-G and AAV8-Flex(FRT)-TVA-mCherry (1:1; 1.82×10^13^ vg/ml and 1.39×10^13^ vg/ml; Stanford gene vector and viral Core) and 4 weeks later 800nl EnvA G-Deleted Rabies-eGFP (Salk viral core). We also unilaterally injected 80-100nl of AAVDJ-Ef1a-fDIO-EYFP (2.1×10^12^ vg/ml; UNC vector core) into the MeA of Dbx1^cre+/^-;FlpO^+/-^ mice for anterograde tracing experiments. For Chr2-assisted circuit mapping, we injected 150nl of AAV2-Flex-GFP (3.7×10^12^ vg/ml; UNC vector core) unilaterally into the MeA of Foxp2^cre+/-^ mice and 40-200nl AAV9-hSyn-ChrimsonR-tdTomato (5.5 × 10^12^ vg/ml; Addgene) unilaterally into the olfactory bulb (AP: 4.45mm, ML= 0.25mm, DV: −1.55mm) of Foxp2^cre+/-^ and Dbx1^cre+/-^;Ai6^+/-^ mice. EnvA G-deleted Rabies virus titers were >1.00×10^8^transforming units per ml.

During surgery, adult male mice were anaesthetized with isoflurane (2%) and then placed in a stereotaxic apparatus (Kopf Instruments). For fiber photometry recordings in juvenile mice, P11 pups were anesthetized with isoflurane (2%) and placed in a stereotaxic apparatus modified with a neonatal anesthesia head holder and zygoma ear cups (Kopf Instruments). The virus or tracer was then delivered into the target region of interest in pups or adults through a glass capillary by using a nanoinjector (World Precision Instruments). For fiber photometry experiments in adults, a 400μm optical fiber (Thorlabs, FT400EMT) attached to a ceramic ferrule (Thorlabs, CF440-10) was placed 0.3mm dorsal to the viral injection site and cemented with adhesive dental cement (C&B metabond, S380). A 3D printed head-fix ring was also secured with cement to the skull. For juvenile experiments, juveniles at P24 were implanted with the optical fiber in the MeA (AP: −0.7mm, ML= 2.03mm, DV: −4.75mm) but the head-fix ring was not utilized. Histology analysis was performed for all animals and only animals with correct virus expression and fiber placement were used for final analysis.

### Behavioral assays and analysis

Behavior was recorded by two synchronized top and side cameras (Basler, acA640-100gm) at 25 frames/second and a digital video recording software (Streampix 5, Norpix) in a dark-room with infrared lights. Behaviors were manually annotated on a frame-by-frame basis by using a custom Matlab function named ‘BehaviorAnnotator’ (https://github.com/pdollar/toolbox).

For male-male interactions we annotated investigation, groom, mount, and attack. For fiber photometry analysis, investigation and groom have been combined as ‘investigation’. For male-female interactions, we recorded investigation, mount, intromission and ejaculation. For male-pup interactions, we recorded investigation, groom, carry and infanticide. ‘Investigation’ was considered as nose-contact to any body part of the target mouse. ‘Groom’ was classified when a mouse has its front paws holding the back or face of the target mouse and is licking either face or back. ‘Attack’ was determined as a series of actions by which the male mouse lounged, bite, chased and pushed the target mouse. ‘Mount’ was defined as a series of fast movements by which the male mouse placed its front paws on the target mouse and positioned itself on top of the target mouse. ‘Intromission’ was annotated as rhythmic deep thrusts following mount. ‘Ejaculation’ was considered when the male stopped deep thrusting and froze in place for several seconds while strongly holding the target female mouse and then slumping to the side. ‘Carry’ involved the male mouse grabbing the pup with mouth, lifting and dropping it off at another location in the cage. ‘Infanticide’ was considered as biting the pup that result in tissue damage. For chemogenetic analysis, pup investigation and groom, were combined as ‘pup investigation’.

### Fiber photometry

Foxp2^cre+/-^ and Dbx1^cre+/-^;FlpO^+/-^ male mice aged 2-8 months were used for adult fiber photometry recordings. Foxp2^cre^ male mice starting at age P25 were used for juvenile fiber photometry experiments. For adult head-fixed experiments, the mice were naïve and did not have had any interactions with other conspecifics outside of their littermates. The recording mouse was head-fixed using a 3D printed head-ring and placed on a 3D printed wheel ^56^. Mice were trained on the wheel for a minimum of three days for at least 10 minutes each day. Each stimulus was presented 5 times for 10 sec with a 50 sec interval in between presentations and a minimum of 5 min break in between different stimuli. Male and receptive female stimulus mice were anaesthetized with ketamine (100mg/kg) and xylazine (10mg/kg).

Fiber photometry was performed as previously described ^46,57,58^. To analyze changes in Ca^2+^ activity, Matlab function ‘msbackadj’, with a moving window of 25% of the total recording time, was utilized to obtain the instantaneous baseline signal (F_baseline_). The instantaneous ΔF/F was calculated as (F_raw_ –F_baseline_)/F_baseline_). The z-score of the ΔF/F (Fz) was obtained by using the Matlab function ‘zscore’ for the whole trace. The peri-event histograms (PETHs) were calculated by aligning the Fz of each trial to either the onset or offset of each behavior. In recordings of head-fix naïve male mice (**Fig. 2),** the Fz peak was calculated by obtaining the average of the maximum value during stimulus presentation. The male preference index (PI) was calculated as (Z_investigate male_ – 0.5 × (Z_investigate female_ + Z_investigate pup_))/( Z_investigate male_ + 0.5 × |Z_investigate female_ + Z_investigate pup_|); The female PI was calculated as (Z_investigate female_ – 0.5 × (Z_investigate male_ + Z_investigate pup_))/( Z_investigate female_ + 0.5 × |Z_investigate male_ + Z_investigate pup_|); The pup PI was calculated as (Z_investigate pup_ – 0.5 × (Z_investigate male_ + Z_investigate female_))/( Z_investigate pup_ + 0.5 ×

|Z_investigate female_ + Z_investigate male_|).

When recording from freely moving naïve mice (**Figs. 3-5**), a receptive female, an adult male mouse and pup were introduced into the cage for 10 mins each except the pup (P1-7) which was introduced for 5 mins. For freely-moving experienced male mice, a pup was introduced into the resident’s cage for 5 mins, the male intruder was placed in the cage for a minimum of 10 mins until the recording mice elicited >6 attacks, without exceeding a total of 1 hour in the cage. A receptive female was introduced until 5 mins after the recording mouse ejaculated. The response elicited during a behavior was calculated as the average Fz during that behavior, while the Fz peak during introduction was calculated as the peak Fz during the first 100 sec after intruder introduction. The male, female and pup PIs were calculated as aforementioned for head-fixed mice. The introduction male, female and pup PIs were calculated using the average Fz during the first 100 sec of stimulus introduction.

When comparing naïve freely-moving and experienced male mice responses, the latency to respond was calculated as the time lapse from behavior onset to when the response reaches Z >= 1. The ‘percent of trials to respond’ was calculated as the percentage of trials that reached Z >= 1. ‘Sniff per trial(s)’ was calculated as the average duration of all male investigation trials. Heatmaps were constructed as F_Z_ – Fz at time 0 for each trial.

### Chemogenetic mediated activation and silencing

For chemogenetic activation experiments, experimental male mice were naïve and had no prior social experience except their littermates. On day 1, male mice were i.p. injected with saline and 30 min after injection, video recordings started. After a 5 min baseline period, a pup intruder was placed into the cage for 5 mins, followed by a 10 min presentation of an adult male, and a 10 min presentation of a receptive female, with 5 min breaks in between stimulus presentation. On day 2, male mice were i.p. injected with 1mg/kg of CNO (Sigma, C0832) and pup, adult male and receptive female were introduced as in day 1.

For chemogenetic silencing experiments, experimental male mice were trained to attack by introducing an adult male mouse daily for 10-30mins minutes/day until they could reliably attack within a 10 min period. Mice were then i.p. injected with saline or CNO (1mg/kg) on interleaved days for two rounds. Thirty minutes after injection, behavioral recordings started and after a 5 min baseline period, an adult male or a receptive female was introduced into the cage for 10 mins each with a 5 min break. Animals with correct bilateral histology were included for analysis.

### Animal body tracking

The velocity (pixels/frame) of each animal after 30 mins of saline or CNO i.p. injection was obtained during the first 5 mins of the chemogenetic assay prior to introduction of any stimulus. The location of each animal was tracked using the top-view camera recordings and analyzed using a custom-written Matlab GUI and code (https://github.com/pdollar/toolbox) ^39^.

### Immunohistochemistry and imaging analysis

Mice were anesthetized and perfused with 1x PBS followed by 20ml 4% PFA. Brains were fixed in 4% PFA for 6-12 hrs at 4°C and dehydrated in 15% sucrose overnight. Brains were embedded in O.C.T. compound (Sakura, 4583) and cut in 50µm sections using a cryostat (Leica CM1950). Every third section was used for immunohistochemistry. Free floating sections were incubated with primary antibody in PBST (0.3% Triton X in PBS) and blocked in 10% normal donkey serum (Jackson ImmunoResearch, 017-000-121) at room temperature in a shaker overnight. The brain sections were then washed 5x in PBST for 10 mins and placed in secondary antibody in PBST and blocked in 10% normal donkey serum for 4 hours at room temperature or overnight at 4°C degrees. Brain sections were then washed 5x in PBST for 10 mins, mounted (Fisher Scientific, 12-550-15) and cover-slipped using fluoromount mounting media with DAPI (ThermoFisher, 00-4959-52). Primary antibodies used were rabbit anti-Foxp2 (1:500, abcam ab16046), rat anti-GFP (1:1000, Nacalai 04404-84), and rabbit anti-mCherry (1:1000, TaKaRa Living Colors DsRed Polyclonal Ab 632496). Secondary antisera used were donkey anti-rat Alexa 488 (1:300; Jackson ImmunoResearch 712-545-150), and donkey anti-rabbit Cy3 (1:1000, Jackson ImmunoResearch 711-165-152). Sections were imaged using a slide scanner (Olympus, VS120) or a confocal microscope (Zeiss LSM 800). Brain sections were identified based on the Allen Mouse Brain Atlas and counted manually using Adobe Photoshop. Cells stained with DAPI were counted using the ImageJ software to automatically count these cells using the ‘analyze particles’ feature and manually corrected.

### Monosynaptic-retrograde rabies input mapping

To determine the inputs to MeA^Foxp2^ and MeA^Dbx1^ cells we injected adult male mice with Cre or Flp dependent AAV-G and AAV-TVA-mCherry viruses and 4 weeks later with EnvA G-Deleted Rabies-eGFP. After 7 days, mice were perfused and every one in three brain sections were collected (50µm thickness sections). Starter cells were considered TVA-mCherry and Rabies-eGFP double positive. Upstream Rabies-eGFP cells were then counted using the ImageJ software. Due to close proximity with the MeA starter cell location, the LH, anterior MeA and AAA were excluded from analysis. Brains with more than 70% of starter cells in the MeA were considered for further analysis. Regions with more than 2% of total inputs to MeA^Foxp2^ and MeA^Dbx1^ cells were included in **Fig. 6**.

### Output axonal projection mapping

To determine the projection patterns of MeA^Foxp2^ and MeA^Dbx1^ cells, every one in three brain sections were collected (50µm thickness). A box area encompassing each region of interest was selected and average pixel intensity was obtained using Adobe Photoshop and calculated as I_raw_. On the same image, a box area of the same size but on the contralateral side with no terminals was used to calculate the background intensity as I_background_. The I_signal_ was obtained by subtracting I_background_ from I_raw_ and then normalizing the value by the maximum I_signal_ across all brain regions for each animal (I_norm_) ^57^. The average I_norm_ was then calculated for all animals to obtain the average axonal projection intensity for each terminal field. Animals with more than 65% of starter cells in the MeA were considered for analysis. Regions with more than 0.2 normalized intensity were included in **Fig. 8**. The LH and anterior MeA were excluded from analysis due to close proximity to the starter cells.

### Brain slice electrophysiology

For AOB to MeA circuit mapping experiments, we injected AAV2-Flex-eGFP and AAV9-hSyn-ChrimsonR-tdTomato into the MeA and the AOB, respectively, of Foxp2^cre+/-^ male mice; or AAV9-hSyn-ChrimsonR-tdTomato into the AOB of Dbx1^cre+/-^Ai6^+/-^ male mice. Whole cell patch-clamp recordings were performed on MeA slices from all mice.

Mice were anesthetized with isoflurane, and brains were removed and submerged in ice-cold cutting solution containing (in mM): 110 choline chloride, 25 NaHCO3, 2.5 KCl, 7 MgCl2, 0.5 CaCl2, 1.25 NaH2PO4, 25 glucose, 11.6 ascorbic acid and 3.1 pyruvic acid. Coronal sections of 275 um were cut on a Leica VT1200s vibratome and incubated in artificial cerebral spinal fluid (ACSF) containing (in mM): 125 NaCl, 2.5 KCl, 1.25 NaH2PO4, 25 NaHCO3, 1 MgCl2, 2 CaCl2 and 11 glucoses at 34°C for 30 min and then transferred to room temperature for cell recovery until the start of recording. Whole-cell voltage-clamp recordings were performed with micropipettes filled with intracellular solution containing (in mM): 135 CsMeSO3, 10 HEPES, 1 EGTA, 3.3 QX-314 (chloride salt), 4 Mg-ATP, 0.3 Na-GTP and 8 sodium phosphocreatine (pH 7.3 adjusted with CsOH). Signals were recorded using MultiClamp 700B amplifier, digitized by DigiData1550B with sampling rate 20 kHz (Molecular Devices, USA). Data were analyzed using Clampfit (Molecular Devices) or MATLAB (Mathworks). To activate ChrimsonR-expressing axons, brief pulses of full field illumination (pE-300 white; CoolLED, 605 nm, 1 ms duration, 10 repeats, with 6 s interval) were delivered onto the recorded cell. Optogenetically-evoked EPSCs and IPSCs (oESPSs and oIPSCs) were recorded by holding the membrane potential of recorded neurons at −70 and 0 mV, respectively. ACSF, TTX (1 µM), TTX (1 µM) and 4-AP (100 mM were sequentially used to test if optogenetically evoked responses are monosynaptic. All drugs were pre-applied for 5 min in the slice chamber prior to data acquisition. Latency was measured as the time difference when the current exceeded 1.5 folds of standard deviation of baseline compared to the light onset.

### Data and code availability

Data to support the findings and custom-written data analysis code (Matlab) is available upon reasonable request from the corresponding authors.

### Statistics

All statistical analysis was performed using Matlab or Graphpad Prism software. Statistical analysis performed were two-tailed. Parametric tests, including paired and unpaired t-test and one-way ANOVA, were used if distributions passed Shapiro-Wilk normality test (except one-way ANOVA with missing values, and for sample size ≤4 and two-way ANOVA, in which data normality was assumed, but not tested). If data was not normally distributed, non-parametric tests were used. To determine differences between a group and a hypothetical value, a one sample test was performed, followed by an analysis of multiple p-values using the original FDR method of Benjamini and Hochberg at Q=5%, to correct for multiple comparisons. For comparisons between more than 2 groups, one-way ANOVA or RM one-way ANOVA was performed followed by Tukey’s multiple comparisons test (normally distributed data); Friedman test followed by Dunn’s multiple comparisons test (RM, not normally distributed data); or Kruskal-Wallis test followed by Dunn’s multiple comparisons test (non-matching groups, not normally distributed data). For differences between groups with two independent variables, two-way ANOVA was performed followed by Sidak’s multiple comparisons test. All significant p-values <0.05 were indicated on the figures. *p< 0.05; **p<0.01; ***p<0.001; ****p<0.0001. For detailed statistical analysis, see statistic summary table.

## Acknowledgements

We thank all members of Lin lab, Drs. R. Sullivan, J. Dasen and M. Long for their inputs during the course of this study. We thank Y. Jiang and the Genotyping Core Laboratory of NYU Langone Health for genotyping of the mice for this study. We also thank Dr. Naoshige Uchida for kindly providing the AAV8-fDIO-GCaMP6f virus. We thank Drs. A. Pierani and L. Vigier for providing the Dbx1^cre+/-^ mouse line and for primer sequences for genotyping. We thank Dr. R. Palmiter for providing the Foxp2^cre+/-^ mice. This research was supported by a Leon Levy Neuroscience Fellowship and NIMH K99MH127295 (to J.E.L); NIH grants R01MH101377, 1R01HD092596 and U19NS107616 (D.L.); the Mathers Foundation (D.L.); Dean’s Undergraduate Research Funds (to J.B, G.S. and M.G); Collegiate Research Initiative (to J.B.); R01DA020140, R21MH129995 and the PNC Charitable Trust (J.G.C).

## Author contributions

J.E.L., D.L. and J.G.C. conceived the project. J.E.L. and D.L. designed experiments, analyzed the data and co-wrote the manuscript. D.L. supervised the project. J.E.L. conducted most experiments. L.Y. performed *in vitro* electrophysiology experiments. C.S. assisted with chemogenetic and fiber photometry experiments. J.B. G.S. and M.G. assisted with histology and behavior annotation. N.P. worked on preliminary characterization of axonal projections. J.G.C. provided feedback throughout the course of the study and supervised N.P.

## Extended Data Figures and Legends

**Extended Data Fig 1.**
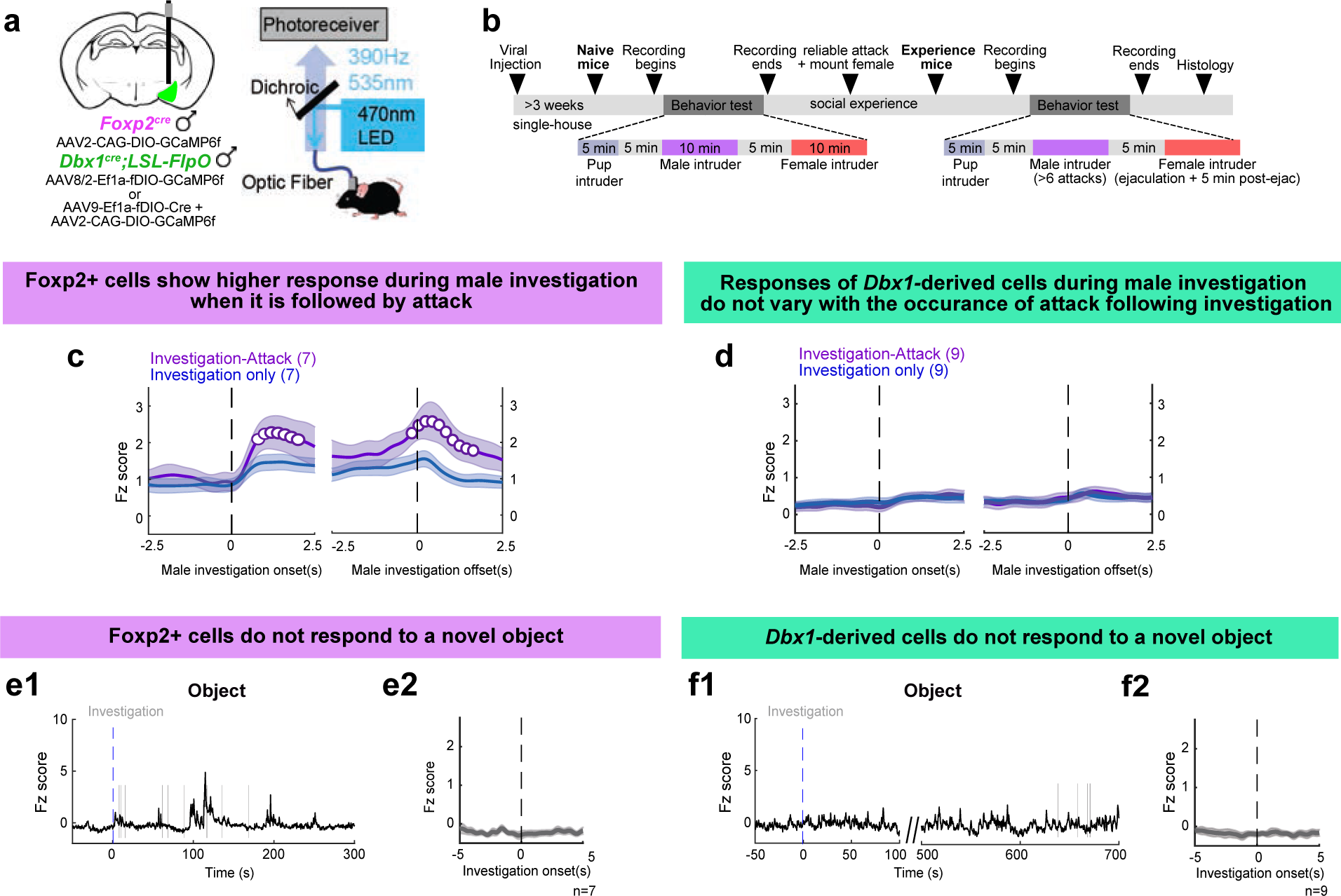
Additional characterization of MeA^Foxp2^ and MeA^Dbx1^ cell responses in experienced male mice, related to Fig. 3. **(a)** Schematic of viral strategy for fiber photometry recordings and fiber photometry setup. **(b)** Experimental timeline for Ca^2+^ recordings in freely-moving naïve and experienced male mice. **(c)** Average PETHs of MeA^Foxp2^ Ca^2+^ signal aligned to the onset (left) and offset (right) of investigation only (blue) and investigation followed by attack (purple). Open circles denote the time period when the investigation-only and investigation-followed-by-attack responses are significantly different (q<0.05). **(d)** Average PETHs of MeA^Dbx1^ cell responses aligned to the onset (left) and offset (right) of investigation only (blue) and investigation followed by attack (purple). The two traces do not differ significantly at any time point. **(e and f)** Representative Ca^2+^ traces (e1, f1) and PETHs (e2, f2) of MeA^Foxp2^ (e) and MeA^Dbx1^ (f) cells during the presentation of a novel object. (c and d) One sample t-test, corrected for multiple comparisons with FDR 0.05. n= number of mice. Data are mean ± S.E.M.

**Extended Data Fig 2.**
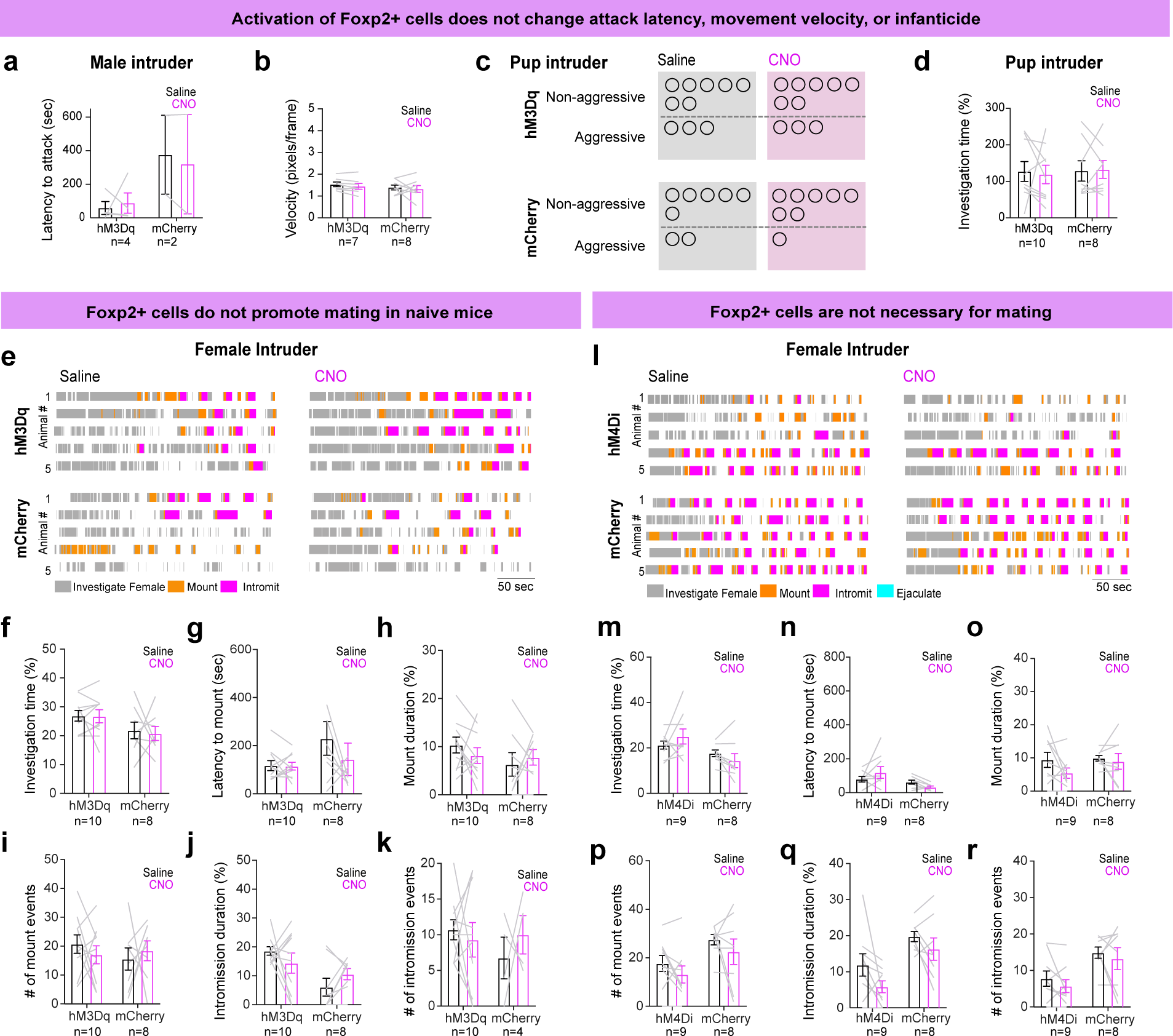
Additional behavioral assays during chemogenetic activation and inactivation of MeA^Foxp2^ cells, related to Fig. 7. **(a)** In Foxp2^hM3Dq^ and Foxp2^mCherry^ male mice, latency to attack a male intruder after CNO injection did not differ from that after saline injection. Only animals that showed attack after both saline and CNO injections were included for this analysis. **(b)** No changes in velocity (pixels/frame) in Foxp2^hM3Dq^ or Foxp2^mCherry^ male mice were observed in a 5 min period 30 min after CNO or saline injection when the test animal was alone in its cage. **(c)** Number of Foxp2^hM3Dq^ or Foxp2^mCherry^ male mice that attacked pups vs. those that did not after saline or CNO injection. Each circle represents one mouse. **(d)** Percentage of time Foxp2^hM3Dq^ or Foxp2^mCherry^ male mice spent investigating the pup after saline or CNO injection. **(e)** Representative raster plots showing the behaviors of 5 Foxp2^hM3Dq^ and 5 Foxp2^mCherry^ mice after i.p. injection of saline or CNO in the presence of a female intruder. **(f-k)** Between CNO-injected and saline-injected days, there is no difference in any parameters related to male sexual behaviors in Foxp2^mCherry^ as well as Foxp2^hM3Dq^ male mice. **(l)** Representative raster plots showing the behaviors of 5 Foxp2^hM4Di^ and 5 Foxp2^mCherry^ mice after i.p. injection of saline or CNO in the presence of a female intruder. **(m-r)** Between CNO-injected and saline-injected days, there is no difference in any parameters related to male sexual behaviors in Foxp2^mCherry^ as well as Foxp2^hM4Di^ male mice. (b, d, f-k, m-r) Two-way repeated measures ANOVA followed by Sidak’s multiple comparisons test; (c) McNemar’s test. n = number of mice. Data are mean ± S.E.M.

**Extended Data Fig 3.**
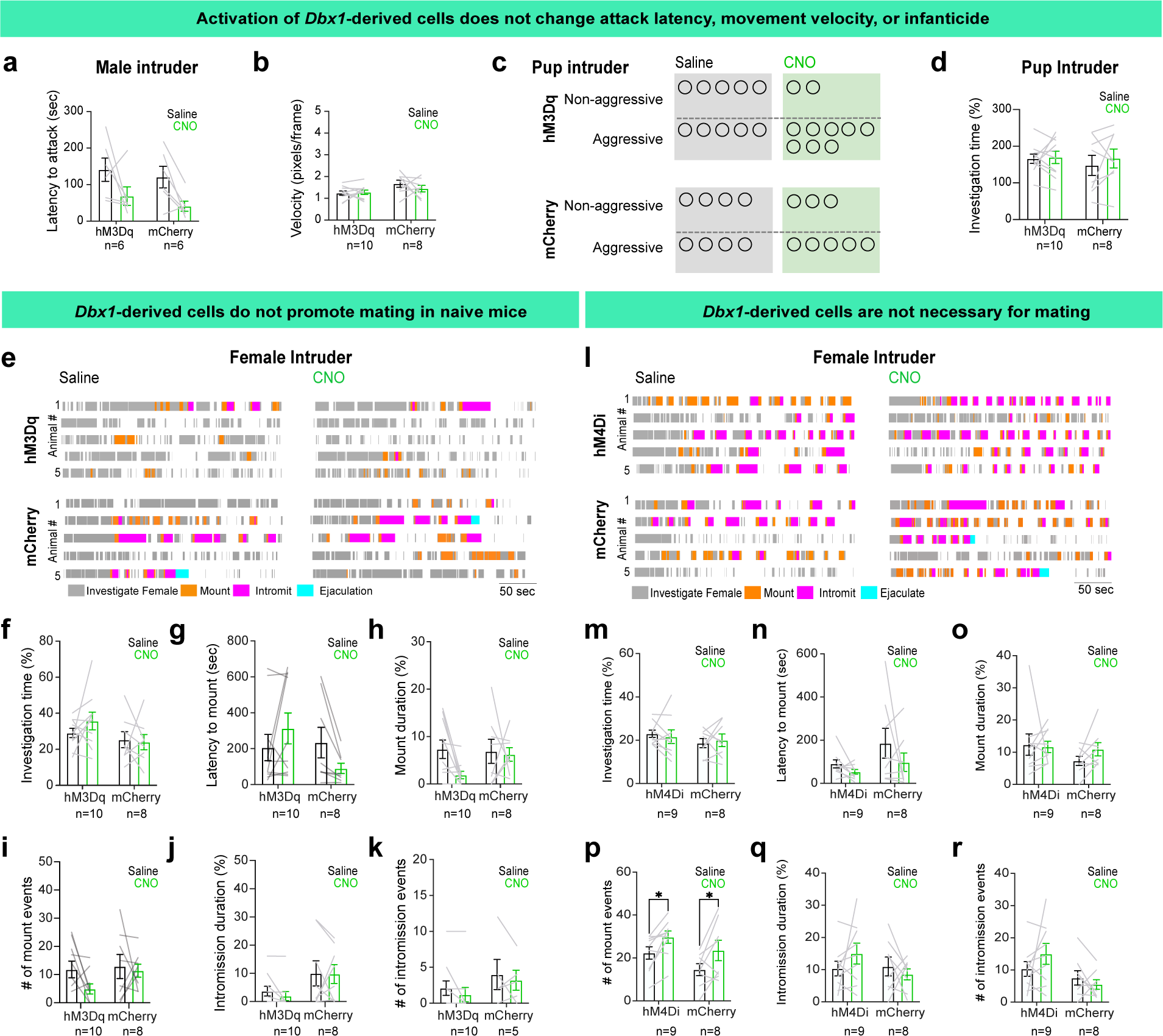
Additional behavioral assays during chemogenetic activation and inactivation of MeA^Dbx1^ cells, related to Fig. 7. **(a)** In Dbx1^hM3Dq^ and Dbx1^mCherry^ male mice, latency to attack a male intruder after CNO injection did not differ from that after saline injection. Only animals that showed attack after both saline and CNO injections were included for this analysis. **(b)** Velocity (pixels/frame) of Dbx1^hM3Dq^ or Dbx1^mCherry^ male mice in a 5 min period after 30 min CNO or saline injection when the test animal was alone in its home cage. **(c)** Number of Dbx1^hM3Dq^ and Dbx1^mCherry^ male mice that attacked pups vs. those that did not after saline or CNO injection. Each circle represents one mouse. **(d)** Percentage of time Dbx1^hM3Dq^ and Dbx1^mCherry^ male mice spent investigating the pup after saline or CNO injection. **(e)** Representative raster plots showing the behaviors of 5 Dbx1^hM3Dq^ and 5 Dbx1^mCherry^ mice after i.p. injection of saline or CNO in the presence of a female intruder. **(f-k)** No difference in male sexual behaviors after CNO injection in comparison to saline injection in Dbx1^mCherry^ nor in Dbx1^hM3Dq^ male mice. **(l)** Representative raster plots showing the behaviors of 5 Dbx1^hM4Di^ and 5 Dbx1^mCherry^ mice after i.p. injection of saline or CNO in the presence of a female intruder. **(m-r)** No difference in male sexual behaviors after CNO injection in comparison to saline injection in Dbx1^mCherry^ nor in Dbx1^hM4Di^ male mice. (a, b, d, f-k, m-r) Two-way repeated measures ANOVA followed by Sidak’s multiple comparisons test; (c) McNemar’s test. n = number of animals. Data are mean ± S.E.M.

**Extended Data Fig 4.**
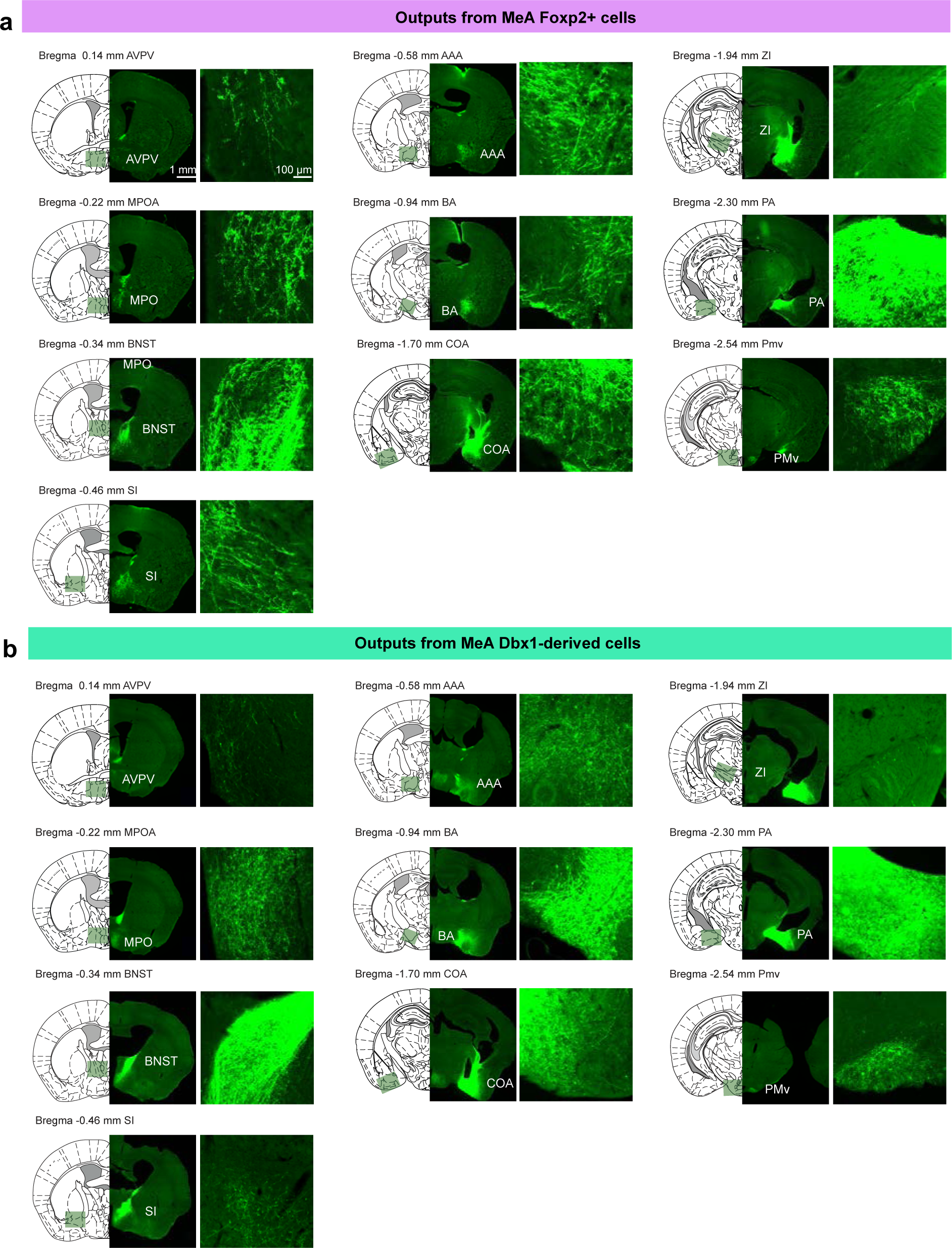
Brain regions downstream of MeA^Foxp2^ and MeA^Dbx1^ cells related to Fig. 8. (a-b) Representative images of 10 brain regions showing the GFP fibers originating from MeA^Foxp2^ (a) and MeA^Dbx1^ (b) cells. The gain of PA and BNST images in (b) was reduced to avoid complete saturation.

