## Supplemental Figures for "Hardwired to attack: Transcriptionally defined amygdala subpopulations play distinct roles in innate social behaviors"

### Extended Data Figures and Legends

### Extended Data Figure 1, related to Figure 3

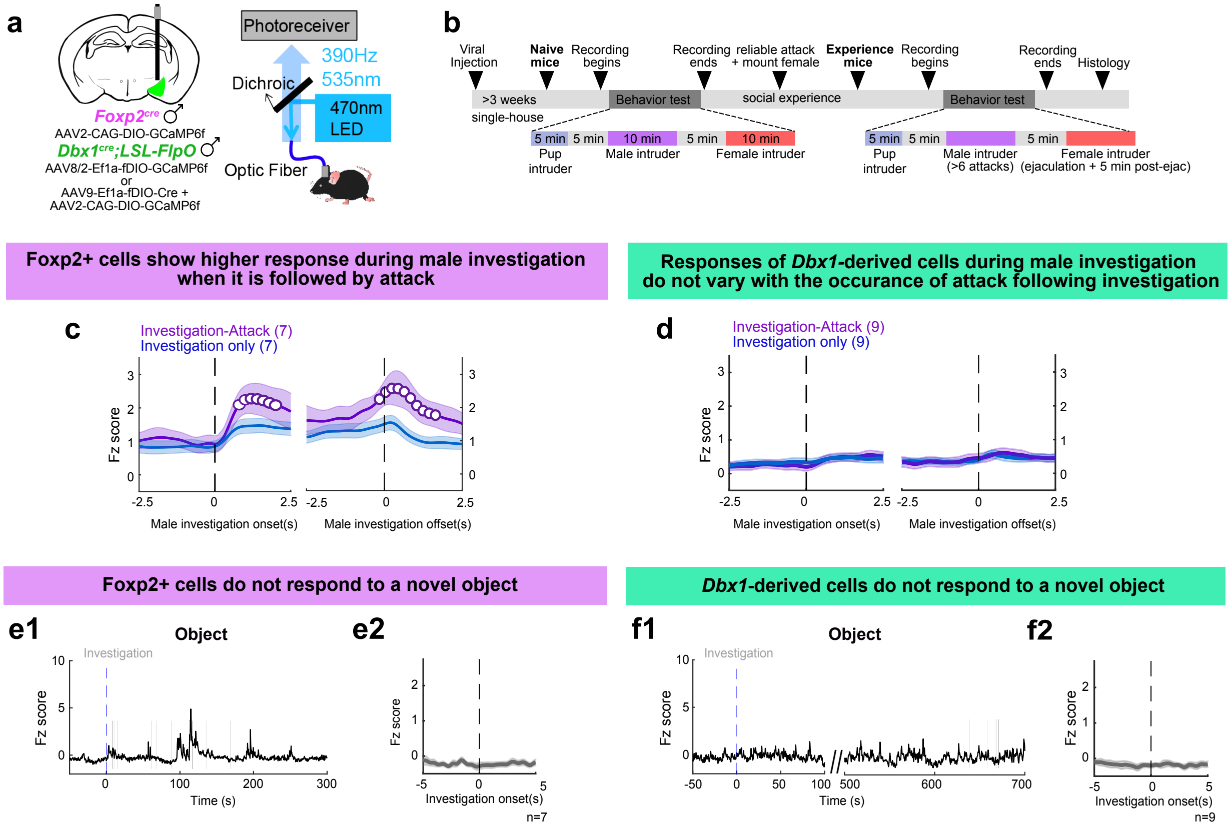


**Extended Data Fig 1. Additional characterization of MeA$^{Foxp2}$ and MeA$^{Dbx1}$ cell responses in experienced male mice, related to Fig. 3.**

**(a)** Schematic of viral strategy for fiber photometry recordings and fiber photometry set-up.

**(b)** Experimental timeline for $Ca^{2+}$ recordings in freely-moving naïve and experienced male mice.

**(c)** Average PETHs of MeA$^{Foxp2}$ $Ca^{2+}$ signal aligned to the onset (left) and offset (right) of investigation only (blue) and investigation followed by attack (purple). Open circles denote the time period when the investigation-only and investigation-followed-by-attack responses are significantly different (q<0.05).

**(d)** Average PETHs of MeA$^{Dbx1}$ cell responses aligned to the onset (left) and offset (right) of investigation only (blue) and investigation followed by attack (purple). The two traces do not differ significantly at any time point.
**(e and f)** Representative $Ca^{2+}$ traces (e1, f1) and PETHs (e2, f2) of MeA$^{Foxp2}$ (e) and MeA$^{Dbx1}$ (f) cells during the presentation of a novel object.

(c and d) One sample t-test, corrected for multiple comparisons with FDR 0.05. n= number of mice. Data are mean ± S.E.M.

### Extended Data Figure 2, related to Figure 7

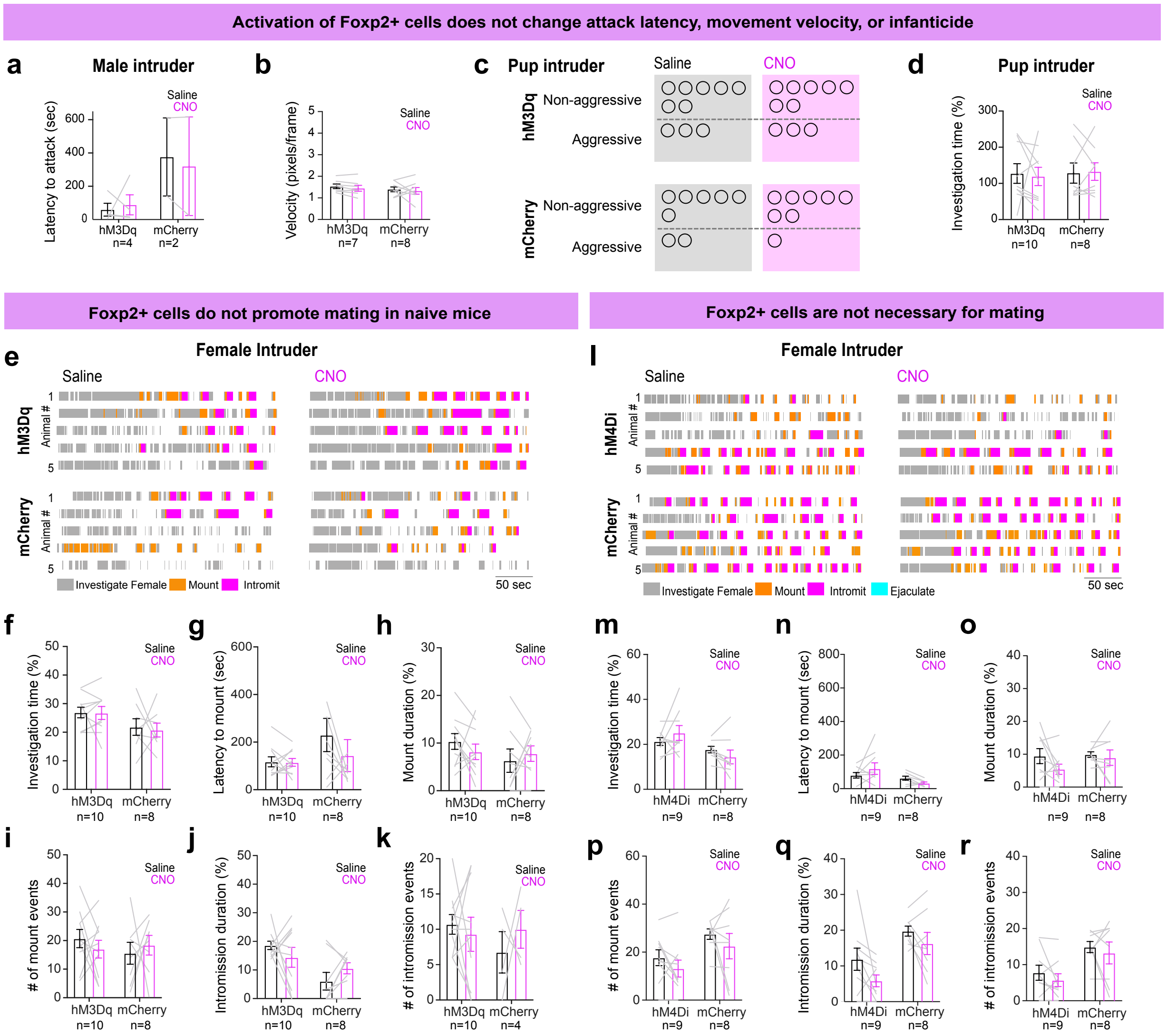


**Extended Data Fig 2. Additional behavioral assays during chemogenetic activation and inactivation of MeA$^{Foxp2}$ cells, related to Fig. 7.**

**(a)** In Foxp2$^{hM3Dq}$ and Foxp2$^{mCherry}$ male mice, latency to attack a male intruder after CNO injection did not differ from that after saline injection. Only animals that showed attack after both saline and CNO injections were included for this analysis.

**(b)** No changes in velocity (pixels/frame) in Foxp2$^{hM3Dq}$ or Foxp2$^{mCherry}$ male mice were observed in a 5 min period 30 min after CNO or saline injection when the test animal was alone in its cage.

**(c)** Number of Foxp2$^{hM3Dq}$ or Foxp2$^{mCherry}$ male mice that attacked pups vs. those that did not after saline or CNO injection. Each circle represents one mouse.
**(d)** Percentage of time Foxp2$^{hM3Dq}$ or Foxp2$^{mCherry}$ male mice spent investigating the pup after saline or CNO injection.
**(e)** Representative raster plots showing the behaviors of 5 Foxp2$^{hM3Dq}$ and 5 Foxp2$^{mCherry}$ mice after i.p. injection of saline or CNO in the presence of a female intruder.
**(f-k)** Between CNO-injected and saline-injected days, there is no difference in any parameters related to male sexual behaviors in Foxp2$^{mCherry}$ as well as Foxp2$^{hM3Dq}$ male mice.
**(l)** Representative raster plots showing the behaviors of 5 Foxp2$^{hM4Di}$ and 5 Foxp2$^{mCherry}$ mice after i.p. injection of saline or CNO in the presence of a female intruder.
**(m-r)** Between CNO-injected and saline-injected days, there is no difference in any parameters related to male sexual behaviors in Foxp2$^{mCherry}$ as well as Foxp2$^{hM4Di}$ male mice.

(b, d, f-k, m-r) Two-way repeated measures ANOVA followed by Sidak's multiple comparisons test; (c) McNemar's test. n = number of mice. Data are mean ± S.E.M.

### Extended Data Figure 3, related to Figure 7

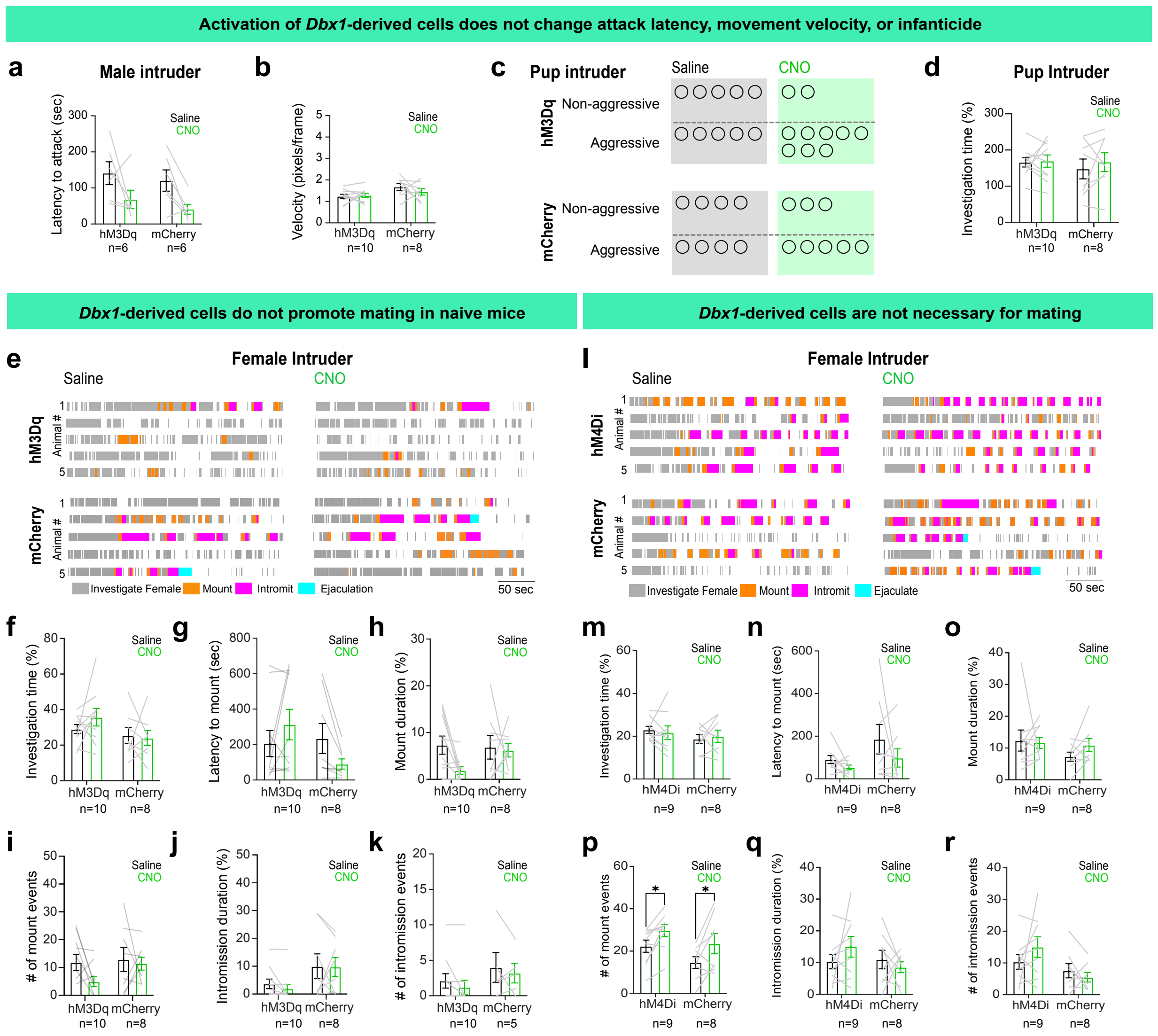


**Extended Data Fig 3. Additional behavioral assays during chemogenetic activation and inactivation of MeA$^{Dbx1}$ cells, related to Fig. 7.**

**(a)** In Dbx1$^{hM3Dq}$ and Dbx1$^{mCherry}$ male mice, latency to attack a male intruder after CNO injection did not differ from that after saline injection. Only animals that showed attack after both saline and CNO injections were included for this analysis.

**(b)** Velocity (pixels/frame) of Dbx1$^{hM3Dq}$ or Dbx1$^{mCherry}$ male mice in a 5 min period after 30 min CNO or saline injection when the test animal was alone in its home cage.

**(c)** Number of Dbx1$^{hM3Dq}$ and Dbx1$^{mCherry}$ male mice that attacked pups vs. those that did not after saline or CNO injection. Each circle represents one mouse.

**(d)** Percentage of time Dbx1$^{hM3Dq}$ and Dbx1$^{mCherry}$ male mice spent investigating the pup after saline or CNO injection.

**(e)** Representative raster plots showing the behaviors of 5 Dbx1$^{hM3Dq}$ and 5 Dbx1$^{mCherry}$ mice after i.p. injection of saline or CNO in the presence of a female intruder.

**(f-k)** No difference in male sexual behaviors after CNO injection in comparison to saline injection in Dbx1$^{mCherry}$ nor in Dbx1$^{hM3Dq}$ male mice.

**(l)** Representative raster plots showing the behaviors of 5 Dbx1$^{hM4Di}$ and 5 Dbx1$^{mCherry}$ mice after i.p. injection of saline or CNO in the presence of a female intruder.

**(m-r)** No difference in male sexual behaviors after CNO injection in comparison to saline injection in Dbx1$^{mCherry}$ nor in Dbx1$^{hM4Di}$ male mice.

(a, b, d, f-k, m-r) Two-way repeated measures ANOVA followed by Sidak's multiple comparisons test; (c) McNemar's test. n = number of animals. Data are mean ± S.E.M.

### Extended Data Figure 4, related to Figure 8

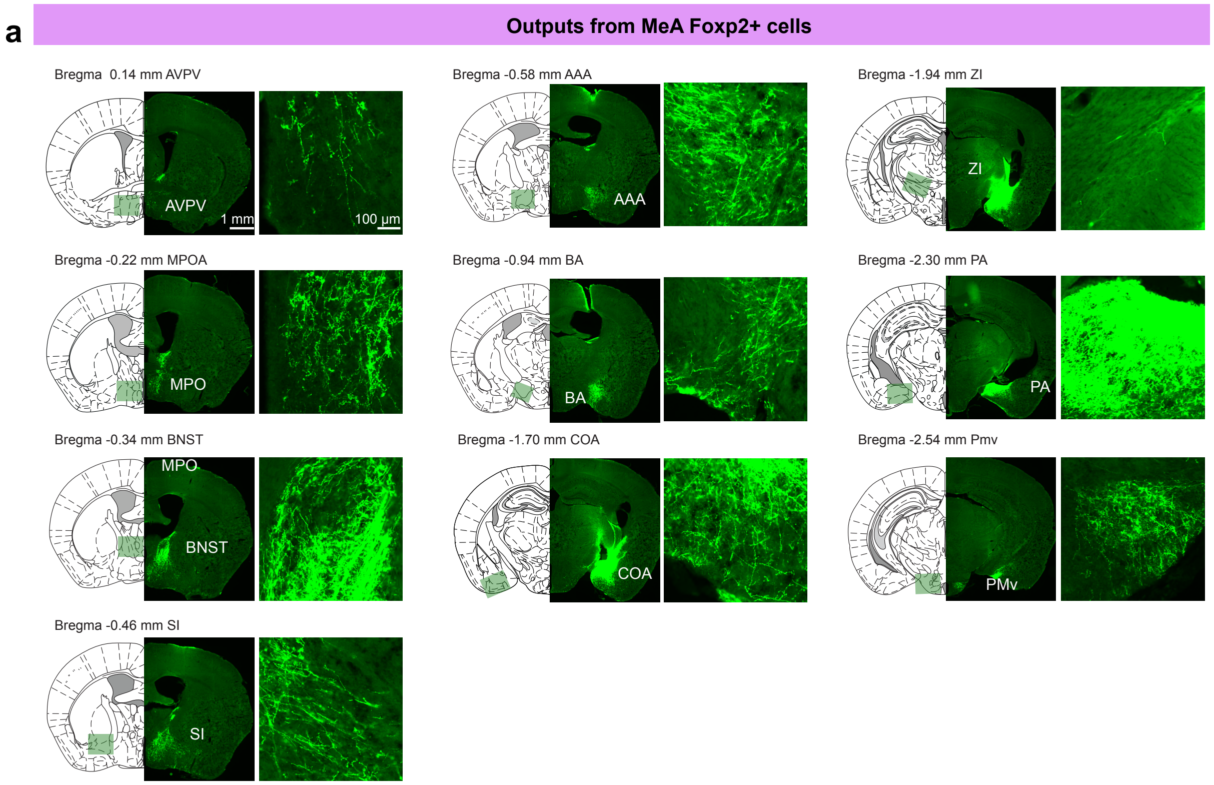


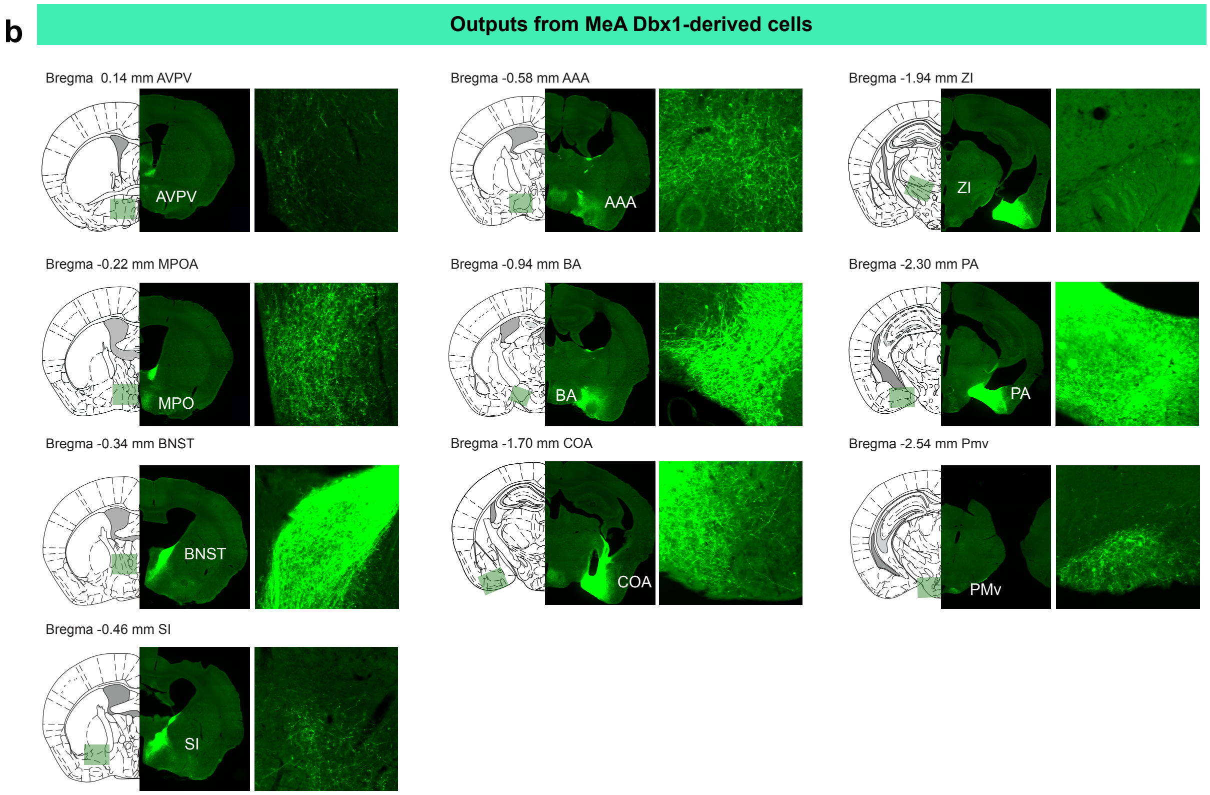

**Extended Data Fig 4. Brain regions downstream of $MeA^{Foxp2}$ and $MeA^{Dbx1}$ cells related to Fig. 8.**

(a-b) Representative images of 10 brain regions showing the GFP fibers originating from $MeA^{Foxp2}$ (a) and $MeA^{Dbx1}$ (b) cells. The gain of PA and BNST images in (b) was reduced to avoid complete saturation.
